# A Conserved SIFamide Circuit Functions as the Insect Vagus Nerve

**DOI:** 10.64898/2026.08.23.746484

**Authors:** Yanying Sun, Yanan Wei, Yutong Song, Wenjing Li, Kyle Wong, Jie Chen, Shun-Fan Wu, Tianmu Zhang, Woo Jae Kim

## Abstract

The insect functional homolog of the mammalian vagus nerve has remained elusive. Here, we identify the SIFamide (SIFa) peptidergic circuit as this homolog in *Drosophila melanogaster*. SIFa neurons project the longest axons from the brain to the hindgut ampulla, forming synapses with peripheral SIFa-receptor (SIFaR) neurons originating in the abdominal ganglion. This circuit is functionally plastic: sexual experience enhances synaptic strength at the hindgut, increasing excretion while reducing mating duration to t**o** favor energy conservation and metabolic recovery. Mechanistically, this shift involves octopaminergic/tyraminergic gating and SIFa-mediated modulation of Adipokinetic Hormone (AKH) from the corpora cardiaca. We further uncover a feedback loop in which SIFaR/Allatostatin-A (AstA) neurons project back to the brain, signaling hindgut physiological status to SIFa neurons via AstA-R1. This hindgut projection is conserved across diverse insect orders. Thus, the SIFa-SIFaR circuit constitutes an evolutionarily conserved “insect vagus nerve” that integrates metabolic state and reproductive history to optimize survival strategies.

## INTRODUCTION

Survival in a fluctuating environment requires the precise coordination of external behavior with internal physiological states. Organisms must continuously balance competing drives—most notably, the trade-off between energy acquisition (foraging and feeding) and genetic propagation (mating and reproduction) ^1^. In mammals, this integration is largely mediated by the vagus nerve, a primary component of the parasympathetic nervous system. The vagus nerve functions as a bidirectional superhighway, transmitting visceral information from the gut to the brain (interoception) and delivering efferent signals from the brain to regulate cardiac activity, digestion, and metabolic homeostasis ^2–6^. While the fundamental principles of energy homeostasis are conserved across phyla, the functional equivalent of the vagus nerve in insects has remained elusive.

In insects, the regulation of feeding and metabolism is known to involve distributed neuroendocrine networks, such as the stomatogastric nervous system and various localized ganglia ^7–13^. Neuropeptides like Allatostatin-A (AstA), Insulin-like peptides (Ilps), and Adipokinetic hormone (AKH) play established roles in nutrient sensing and satiety ^11,14–26^. However, a centralized neural substrate that physically connects the higher brain directly to the posterior viscera—and integrates this connection with reproductive state—has not been fully defined. Identifying such a pathway is critical to understanding how insects manage the “decision” to prioritize feeding over mating when metabolic resources are depleted or when reproductive potential is temporarily exhausted.

We focused on SIFamide (SIFa), a strictly conserved neuropeptide in holometabolous insects, as a candidate for this integrative role ^27–31^. SIFa-expressing neurons in the *Drosophila* brain, specifically in the Pars Intercerebralis (PI), are historically characterized as master regulators of sexual behavior ^32–38^. However, the comprehensive morphology of these neurons suggests a function far broader than reproductive control alone. Preliminary observations indicated that SIFa neurites extend well beyond the brain, projecting into the ventral nerve cord (VNC) and interacting with neurohemal organs, hinting at a systemic regulatory capacity reminiscent of the vagus nerve’s broad innervation profile ^28,29,34,39–42^.

In this study, we demonstrate that SIFa neurons form the core of an insect “vagus-like” system. We reveal that SIFa neurons project the longest identified axons in the *Drosophila* nervous system, extending from the brain directly to the hindgut ampulla. There, they form functional synapses with SIFa-receptor (SIFaR) expressing neurons that originate in the abdominal ganglion. We characterize a bidirectional feedback loop wherein these peripheral SIFaR neurons, which co-express AstA, project axons back to the brain (SOG/PWR regions) to modulate SIFa neuronal activity. Furthermore, we show that this circuit is functionally plastic: sexual satiety in males strengthens the brain-hindgut synaptic connection via an Tyraminergic (TA) or Octopaminergic (OA) gating mechanism. This potentiation shifts the animal’s behavioral state from mating investment to energy conservation and metabolic restoration, mediated centrally by the modulation of AKH in the Corpora Cardiaca (CC) and peripherally by altered hindgut physiology. Finally, we provide evidence that this direct brain-hindgut projection is conserved in other insect orders, including *Hemiptera* (*Nilaparvata lugens,* brown planthopper), *Lepidoptera* (*Spodoptera frugiperda,* fall armyworm), and *Hymenoptera* (*Messor structor*, harvest ant). Collectively, our findings define the SIFa-SIFaR circuit as a functional evolutionary homolog of the vagus nerve, essential for coupling reproductive history with metabolic homeostasis.

## RESULTS

### SIFa-Mediated Coordination of the Gut-Brain Axis in Post-Mating Males

To identify a functional homolog of the vagus nerve in *Drosophila*, we first established specific behavioral and physiological criteria. In mammals, the vagus nerve serves as the primary conduit for the “rest and digest” system, bidirectionally coupling the central nervous system with visceral organs to coordinate energy homeostasis ^2,4,5,43,44^. A true insect homolog must therefore do more than simply regulate a single organ; it must act as a master switch that rebalances the organism’s internal state, specifically mediating the trade-off between energy expenditure (e.g., reproduction) and energy acquisition (e.g., feeding and digestion) ^2,45–48^.

We identified the post-mating male *Drosophila* as the ideal biological model to test for this systemic integration. While male mating is often viewed merely as a reproductive endpoint, recent syntheses demonstrate that successful copulation triggers a “profound metamorphosis” in the male, initiating an active, coordinated, and adaptive biological program ^49–55^. This state represents a fundamental shift in “ejaculate economics,” where the male must manage finite seminal resources against future reproductive opportunities ^56–63^.

Crucially, this post-mating transformation is a multi-system event that mirrors the homeostatic regulation of the vagus nerve. It involves a “deep physiological reconfiguration” characterized by metabolic costs and the mobilization of energy reserves alongside a “hierarchical behavioral shift” where high-energy drives like courtship are suppressed in favor of recovery and strategic resource management. The post-mating male essentially transitions from a state of high reproductive investment to a state of metabolic replenishment—a “rest and digest” phase that requires precise coordination between the brain and the gut ^12,23,64–76^.

To experimentally dissect the role of the SIFa circuit in this transition, we selected two complementary phenotypic readouts acting as proxies for the competing drives of the vagus axis. We utilized the Excretion Quantification (EX-Q) assay to monitor hindgut physiology; because the post-mating state compels the male to offset the significant “metabolic price” of reproduction, changes in excretion and gut throughput serve as a direct readout of the visceral regulation required for energy balance ^77^. We coupled this with an analysis of the Shorter-Mating-Duration (SMD) response, a behavioral metric of reproductive investment ^78,79^. This behavior is not a simple reflex but a “memory-guided, sensory-gated, and metabolically-tuned” strategy that represents a calculated reduction in reproductive time-investment by experienced males to conserve their finite ejaculate resources ^80–82^.

By assessing how the SIFa circuit modulates these two distinct biological outputs— one visceral (EX-Q) and one behavioral (SMD)—we aimed to determine if SIFa neurons function as the neural substrate linking metabolic status in the gut with reproductive decision-making in the brain ^36–38^. Consistent with this hypothesis, we found that males subjected to excessive mating displayed a dramatic increase in excretion levels (Fig. 1A), signaling a shift toward metabolic processing. This surge in visceral activity was accompanied by a concurrent reduction in mating duration (Fig. 1B), recapitulating the previously characterized Shorter-Mating-Duration (SMD) phenotype ^78,79^. Crucially, RNAi-mediated knockdown of SIFa abolished both behaviors (Fig. 1C–D). These results demonstrate that the male post-mating state— defined by increased visceral throughput (excretion) and reduced reproductive investment (mating duration)—is actively coordinated via SIFa-mediated signaling (Fig. 1E).

**Figure 1.**
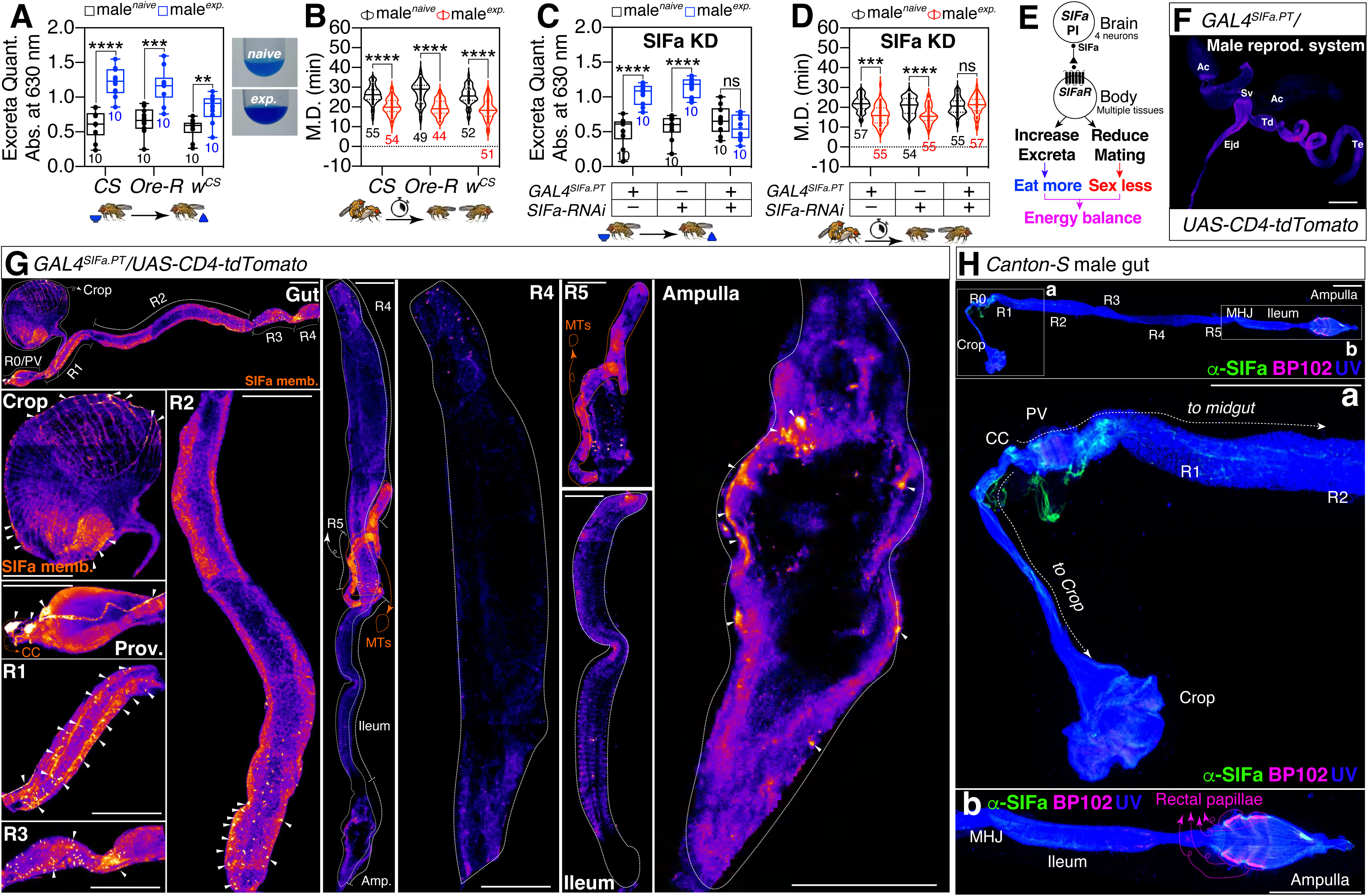
SIFa neurons coordinate the post-mating behavioral-physiological shift and project long axons directly to the hindgut. (A) 24-h sucrose intake of males measured by EX-Q assay of *Canton-S, Oregon-R,* and *Cantonized*-*white* (wCS) male flies on yeast-sugar medium. See the **MATERIALS AND METHODS** for a detailed description of the EX-Q assay used in this study. (B) SMD assays for *Canton-S, Oregon-R,* and *Cantonized*-*white* male flies. In the mating duration (MD) assays, black violin plot denote males that were sexually naïve flies, whereas red violin plot denote males that were sexually experienced. The numerical values beneath the violin plots indicate the count of male flies that mated successfully. The mean value and standard error are labeled within the violin plot (black lines and red lines). M.D represent mating duration. (See **MATERIALS AND METHODS**). Asterisks represent significant differences, as revealed by the unpaired Student’s t test, and ns represents non-significant differences (*\*p<0.05, **p<0.01, ***p< 0.001, ****p< 0.0001*). Consequently, data points on graphs marked with asterisks indicate that SMD behaviors remain unaltered or within normal parameters, whereas those labeled with ‘ns’ signify that SMD behaviors have been perturbed due to mutations or genetic alterations in the respective strains. For detailed methods, see the **MATERIALS AND METHODS** for a detailed description of the MD assays used in this study. In the framework of our investigation, the routine application of internal controls is employed for the vast majority of experimental procedures, as delineated in the “**Mating Duration Assay**” and “**Statistical Tests**” subsections of the **MATERIALS AND METHODS** section. The identical analytical approach employed for the MD assays is maintained for the subsequent data presented. (C) 24-h sucrose intake of males measured by EX-Q assays of *GAL4^SIFa.PT^*-mediated knockdown of SIFa *via SIFa-RNAi* male flies on yeast-sugar medium. Genetic control assays were performed using heterozygous *SIFa-RNAi*/+ and *GAL4^SIFa.PT^*/+ males. (D) SMD assays of *GAL4^SIFa.PT^*-mediated knockdown of SIFa *via SIFa-RNAi* male flies on yeast-sugar medium. Genetic control assays were performed using heterozygous *SIFa-RNAi*/+ and *GAL4^SIFa.PT^*/+ males. (E) Diagram of how SIFa-SIFaR signaling mediated energy balance through multiple tissues. (F-G) Testis (F), gut (G) of male flies expressing *GAL4^SIFa.PT^* together with *UAS-CD4-tdTomato,* were immunostained with anti-DsRed (red) antibody. The memb. represent cell membrane. For detailed methods, see the **MATERIALS AND METHODS** for a detailed description of the immunostaining procedure used in this study. Scale bars represent 100 μm. (H) Gut of *Canton-S* male flies were immunostained with anti-SIFa (green), BP102 antibody (magenta). The morphology of the entire gut was visualized using UV light. Scale bars represent 100 μm.

To determine whether this SIFa-dependent vagal-like response represents a universal post-mating physiological shift in *Drosophila*, we examined female flies. Strikingly, and in stark contrast to males, RNAi-mediated knockdown of SIFa in females had no effect on the post-mating increase in excretion (Fig. S1A). In females, the post-mating physiological shift is instead governed by independent regulators—most notably the male-derived seminal Sex Peptide (SP) ^62,63,83,84^. Following mating, SP acts on Sex Peptide Receptors (SPR) within the female’s *pickpocket* (*ppk*) and *doublesex* (*dsx*) sensory networks and distinct higher-order processing circuits ^85,86^. to drive broad reproductive and physiological changes ^87–89^, including systemic shifts in gut metabolism, intestinal transit, and excretion ^90,91^. This sexual dimorphism indicates that the SIFa circuit is not a generic mediator of mating-induced gut motility, and the SIFa-dependent vagal-like response linked to sexual experience is not sufficient to drive this excretory function in females. Instead, it operates as a male-specific neural substrate, consistent with the hypothesis that the profound ‘rest and digest’ transition is uniquely critical for males to manage the metabolic costs of ejaculate production and replenish finite reproductive resources ^38^.

To determine if these systemic modulations arise from direct neural regulation, we next mapped the projection targets of SIFa neurons. Using the highly specific *SIFa-PT-GAL4* driver ^35^ to express *tdTomato*, we visualized the comprehensive morphology of SIFa neurites. Imaging revealed extensive innervation throughout the alimentary canal; projections were particularly robust in the foregut and crop. Strikingly, we also observed dense axonal projections extending to the hindgut, specifically concentrating in the ampulla, whereas innervation in the midgut R4 region was comparatively sparse. Notably, despite the profound influence of SIFa signaling on mating behavior, we did not observe direct innervation within the male reproductive organs (Fig. 1F–G). We validated these projection patterns using anti-SIFa antibody staining, which confirmed that SIFa neurons directly innervate both the foregut and hindgut (Fig. 1H). To further characterize the specific topology of this hindgut innervation, we utilized anti-BP102 (a CNS axon marker) staining. This imaging revealed that SIFa axons extend directly to the hindgut ampulla, specifically terminating in close proximity to the four rectal papillae ^92,93^, distinct structures which are clearly demarcated by BP102 (Fig. S1B-D). These anatomical tracings demonstrate that SIFa neurons project from the PI in the brain all the way to the posterior hindgut, constituting what appear to be the longest axonal projections in the *Drosophila* nervous system (Fig. S1E).

### A Bidirectional SIFa-SIFaR Neural Circuit Innervates the Rectal Papillae and Crop to Regulate Visceral Plasticity

Since SIFa signaling is mediated exclusively through the SIFamide receptor (SIFaR) ^36–38,94^, we next investigated the functional requirement of this receptor in the observed behaviors. We found that global knockdown of SIFaR significantly impaired both the post-mating increase in excretion (EX-Q) and the reduction in mating duration (SMD) (Fig. 2A–B). To spatially dissect these functions, we utilized *tsh-GAL80* to inhibit GAL4 activity specifically within the ventral nerve cord (VNC), thereby restricting SIFaR knockdown primarily to the brain while preserving expression in the VNC. Under these conditions, the post-mating excretion phenotype was restored to control levels, whereas the SMD response remained disrupted (Fig. 2C–D). These results suggest a functional dissociation: SIFaR expression within the VNC is critical for the modulation of excretion, whereas the regulation of mating duration appears to be more complex, requiring SIFaR function in both the brain and the VNC as we have previously reported ^36,38^.

**Figure. 2.**
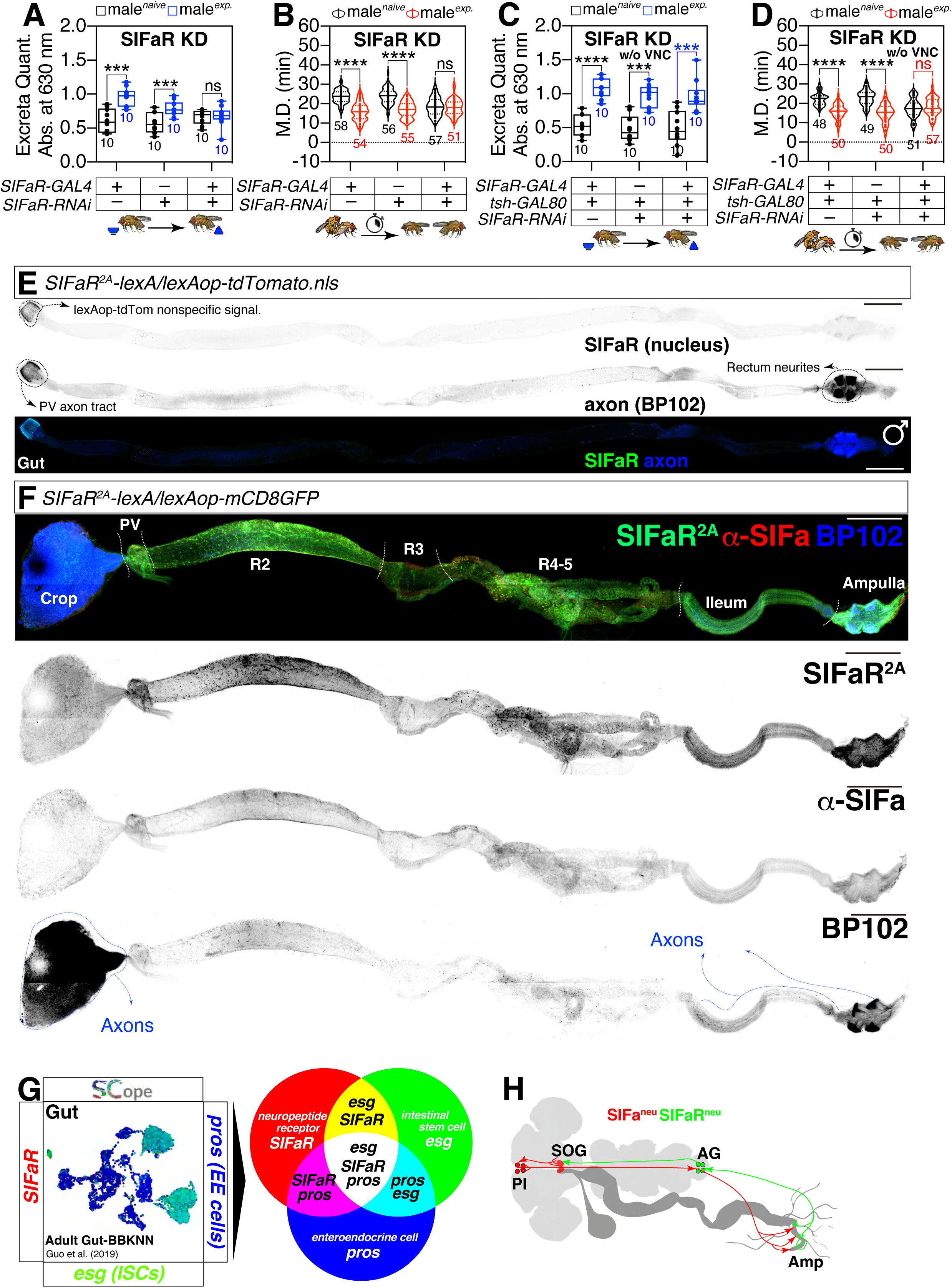
SIFaR functions in abdominal ganglion neurons, rather than intrinsic gut cells, to regulate visceral plasticity. (A) 24-h sucrose intake of males measured by EX-Q assays of *SIFaR-GAL4* mediated knockdown of SIFaR *via SIFaR-RNAi* male flies on yeast-sugar medium. Genetic control assays were performed using heterozygous *SIFaR-RNAi*/+ and *SIFaR-GAL4*/+ males. (B) SMD assays of *SIFaR-GAL4* mediated knockdown of SIFaR *via SIFaR-RNAi* male flies on yeast-sugar medium. Genetic control assays were performed using heterozygous *SIFaR-RNAi*/+ and *SIFaR-GAL4*/+ males. (C) 24-h sucrose intake of males measured by EX-Q assays of *SIFaR-GAL4* mediated knockdown of SIFaR *via SIFaR-RNAi* together with tsh-GAL80 male flies on yeast-sugar medium. Genetic control assays were performed using heterozygous *SIFaR-RNAi, tsh-GAL80*/+ and *SIFaR-GAL4, tsh-GAL80*/+ males. (D) SMD assays of *SIFaR-GAL4* mediated knockdown of SIFaR *via SIFaR-RNAi* together with tsh-GAL80 male flies on yeast-sugar medium. Genetic control assays were performed using heterozygous *SIFaR-RNAi, tsh-GAL80*/+ and *SIFaR-GAL4, tsh-GAL80*/+ males. (E) Gut of male flies expressing *SIFaR^2A^-lexA* together with *lexAop-tdTomato.nls* were immunostained with anti-DsRed (green), and BP102 (blue) antibody. The panels presented as a grey scale is to clearly show the expression patterns of neurons in brain labeled by *SIFaR^2A^*driver. Scale bars represent 100 μm. (F) Gut of male flies expressing *SIFaR^2A^-lexA* together with *lexAop-mCD8GFP* were immunostained with anti-GFP (green), anti-SIFa (red), and BP102 (blue) antibody. The panels presented as a grey scale is to clearly show the expression patterns of neurons in brain labeled by *SIFaR^2A^* driver. Scale bars represent 100 μm. (G) Each tSNE visualization depicts the coexpression patterns of genes, with each color corresponding to the genes listed on the left, right, and bottom of the plot. The tissue name, as referenced on the Fly SCope website is indicated in the upper left corner of the tSNE plot. See the **MATERIALS AND METHODS** for a detailed description of the single-nucleus RNA-sequencing analyses used in this study. (H) Diagram of SIFa-SIFaR circuit from PI region to Ampulla in *Drosophila melanogaster*.

To visualize the circuit architecture, we employed a Chemoconnectome (CCT) knock-in (KI) *SIFaR* line ^95^. Consistent with our hypothesis, we confirmed that *SIFaR* expression is absent within the gut tissue itself (Fig. 2E). Instead, we identified a specific population of *SIFaR*-expressing neurons with cell bodies located in the Abdominal Ganglion (AG) that project axons extensively to both the foregut and hindgut, closely mirroring the innervation pattern of SIFa neurons (Fig. 2F). This lack of intrinsic enteric *SIFaR* expression was independently corroborated by single-cell RNA sequencing data from the FlyCellAtlas (SCope) ^96^, which showed no detectable receptor expression in gut epithelial, muscle, enteroendocrine (EE), or intestinal stem (ISC) cells (Fig. 2G, Fig. S2A–B). These findings indicate that while both SIFa and SIFaR neurites run alongside the gastrointestinal tract, the signaling interface is exclusively neuronal. This architecture—a central command (SIFa) communicating with peripheral ganglia (SIFaR) to regulate visceral function—strongly supports the model of the SIFa-SIFaR axis as a functional homolog of the vagus nerve (Fig. 2H).

A defining characteristic of the vagus nerve is its bidirectional nature, serving as both a sensory and motor pathway to integrate central and visceral states ^97^. To investigate the directionality of information flow within the SIFa-SIFaR axis, we employed the dual-labeling system DenMark/syt-eGFP to distinguish between dendritic inputs and axonal outputs in both neuronal populations ^98^. Consistent with our hypothesis that the SIFa circuit constitutes a bidirectional insect vagus nerve (Fig. 2H), we observed that both SIFa and SIFaR neurons project dendrites and axons throughout the CNS ^36,37^ (Fig. 3, Fig. S3). Crucially, this mixed polarity extends to the periphery: both populations send dendritic and axonal projections to the entire foregut/hindgut region (Fig. 3, Fig. S3), suggesting they are capable of both sending signals to and receiving feedback from these visceral tissues.

**Figure. 3.**
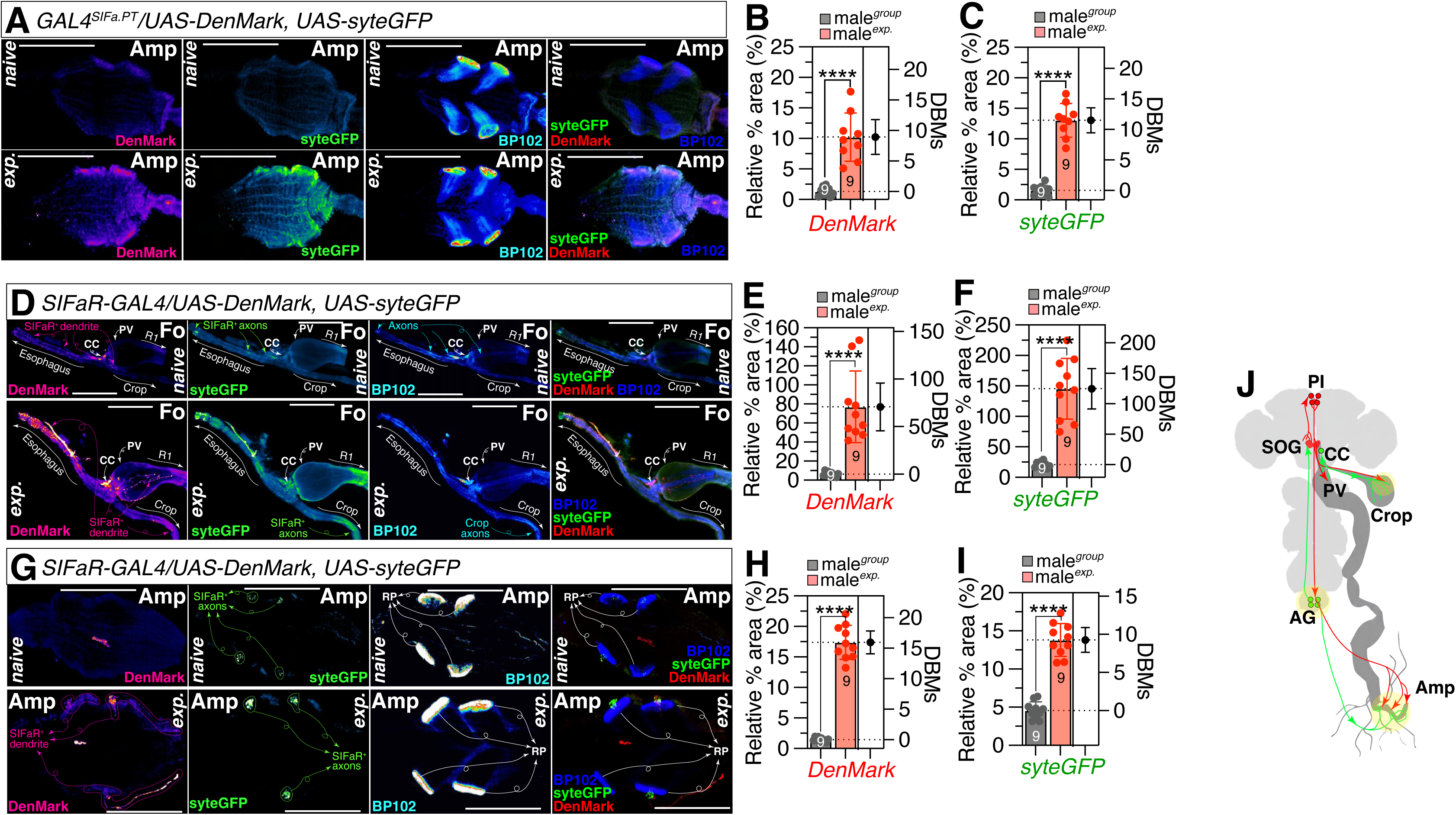
Bidirectional SIFa-SIFaR gastrointestinal projections exhibit mating-induced structural plasticity targeting the rectal papillae. (A) Male flies ampulla expressing *GAL4^SIFa.PT^* together with *UAS-Denmark* and *UAS-syt.eGFP* in naïve (upper) and experienced (lower) conditions. Denmark is pseudo-colored as “fire”. GFP is pseudo-colored as “green fire blue”. BP102 is pseudo-colored as “16 colors”. Scale bars represent 100 μm. (B-C) Quantification of relative intensity value for Denmark and GFP fluorescence in ampulla (two-tailed unpaired *t*-test). In all plots and statistical tests. Data are presented as mean ± s.e.m. ns = not significant (*p>0.05*), *\*p<0.05, **p<0.01, ***p< 0.001, ****p< 0.0001*. Sample sizes (n) are indicated in the figure panels. (D) Male flies foregut expressing *SIFaR-GAL4* together with *UAS-Denmark* and *UAS-syt.eGFP* in naïve (upper) and experienced (lower) conditions. Denmark is pseudo-colored as “fire”. GFP is pseudo-colored as “green fire blue”. BP102 is pseudo-colored as “16 colors”. Scale bars represent 100 μm. (E-F) Quantification of relative intensity value for Denmark and GFP fluorescence in foregut (two-tailed unpaired *t*-test). In all plots and statistical tests. Data are presented as mean ± s.e.m. ns = not significant (*p>0.05*), *\*p<0.05, **p<0.01, ***p< 0.001, ****p< 0.0001*. Sample sizes (n) are indicated in the figure panels. (G) Male flies ampulla expressing *SIFaR-GAL4* together with *UAS-Denmark* and *UAS-syteGFP* in naïve (upper) and experienced (lower) conditions. Denmark is pseudo-colored as “fire”. GFP is pseudo-colored as “green fire blue”. BP102 is pseudo-colored as “16 colors”. Scale bars represent 100 μm. (H-I) Quantification of relative intensity value for Denmark and GFP fluorescence in ampulla (two-tailed unpaired *t*-test). In all plots and statistical tests. Data are presented as mean ± s.e.m. ns = not significant (*p>0.05*), *\*p<0.05, **p<0.01, ***p< 0.001, ****p< 0.0001*. Sample sizes (n) are indicated in the figure panels.

We further observed that this peripheral circuitry exhibits remarkable structural plasticity in response to reproductive state. Using the DenMark/syteGFP system to label dendritic and axonal compartments specifically in SIFa neurons, we found that sexual experience dramatically enhanced both dendritic and axonal signals within the hindgut ampulla (Fig. 3A-C). This indicates that the central SIFa command neurons themselves strengthen their direct connectivity to the hindgut periphery following mating.

Next, we examined the structural plasticity of the downstream SIFaR neurons. Mating also triggered a robust expansion of both dendritic and axonal arbors of SIFaR neurons within the foregut and hindgut regions (Fig. 3D-I). This concurrent potentiation of input and output structures suggests that SIFaR neurons are not static relays; rather, they actively upscale their capacity to both receive sensory feedback and transmit efferent signals when the male enters the mated state.

One of the most extraordinary targets of the SIFa-SIFaR axis is the rectal papillae (RPs), specialized epithelial structures within the hindgut ampulla that are essential for the selective reabsorption of water, ions, and metabolites, thereby maintaining systemic osmoregulation ^92,99^. High-magnification imaging revealed that SIFaR dendrites and axon terminals form a complex, “seed-like” lattice adhering to the dish-shaped surface of the papillae (Fig. S3A). While both input and output structures are present, they exhibit a partially overlapping rather than identical spatial distribution. Detailed tracing further clarified the projection trajectory: SIFaR neurons bypass the midgut-hindgut junction (MHJ) to project directly to the ampulla (Fig. S3B). This peripheral mapping also resolved the connectivity to the reproductive system.

Although central SIFa neurons do not directly innervate the male reproductive organs (Fig. 1F), we observed robust projection of both SIFaR dendrites and axons from AG into these tissues (Fig. S3C). This architecture implies a relay mechanism wherein the central SIFa command modulates reproductive physiology indirectly via these peripheral SIFaR neurons. We next analyzed the synaptic plasticity of the SIFaR-RPs connection in response to sexual experience. Quantitative profiling revealed that the post-mating enhancement of this circuit is driven by a specific structural change: mating significantly increased the total coverage area of synapses projecting to the RPs, without altering the number or size of individual synaptic boutons (Fig. S3D–G).

Finally, we identified the source of the dense SIFaR innervation in the foregut. These projections originate from the Corpora Cardiaca (CC), the insect neurohemal organ analogous to the vertebrate pituitary, which functions as a master regulator of energy homeostasis through the release of metabolic hormones such as Adipokinetic Hormone (AKH) ^100–102^. These SIFaR-positive CC neurons project heavily to the crop, exhibiting a similar overlapping dendritic and axonal pattern to that observed in the rectal papillae (Fig. 3D, Fig. S3H). Collectively, these findings define a spatially segregated organization of the peripheral “vagus” arm: SIFaR neurons in the AG predominantly target the hindgut ampulla and RPs to regulate excretion, while SIFaR neurons in the CC primarily innervate the foregut and crop to modulate metabolic storage (Fig. 3J).

### Synaptic Plasticity and Octopamine-AKH Signaling in the Corpora Cardiaca Coordinate Post-Mating Metabolic Homeostasis

To further investigate the functional architecture of this circuit, we sought to determine if the post-mating expansion of neurites corresponds to increased synaptic connectivity. We utilized t-GRASP (targeted GFP Reconstitution Across Synaptic Partners), a method where split-GFP fragments are tethered to pre- and post-synaptic membranes, reconstituting fluorescence only when specific neurons form functional synapses ^103^. As expected, mated males exhibited a dramatic increase in GRASP signals across the foregut, crop, and hindgut regions, accompanied by elevated levels of SIFa peptide (Fig. 4A–I). These data suggest that the post-mating outgrowth of SIFaR neurites reflects a genuine strengthening of synaptic communication between central SIFa command neurons and peripheral SIFaR targets in the gut.

**Figure. 4.**
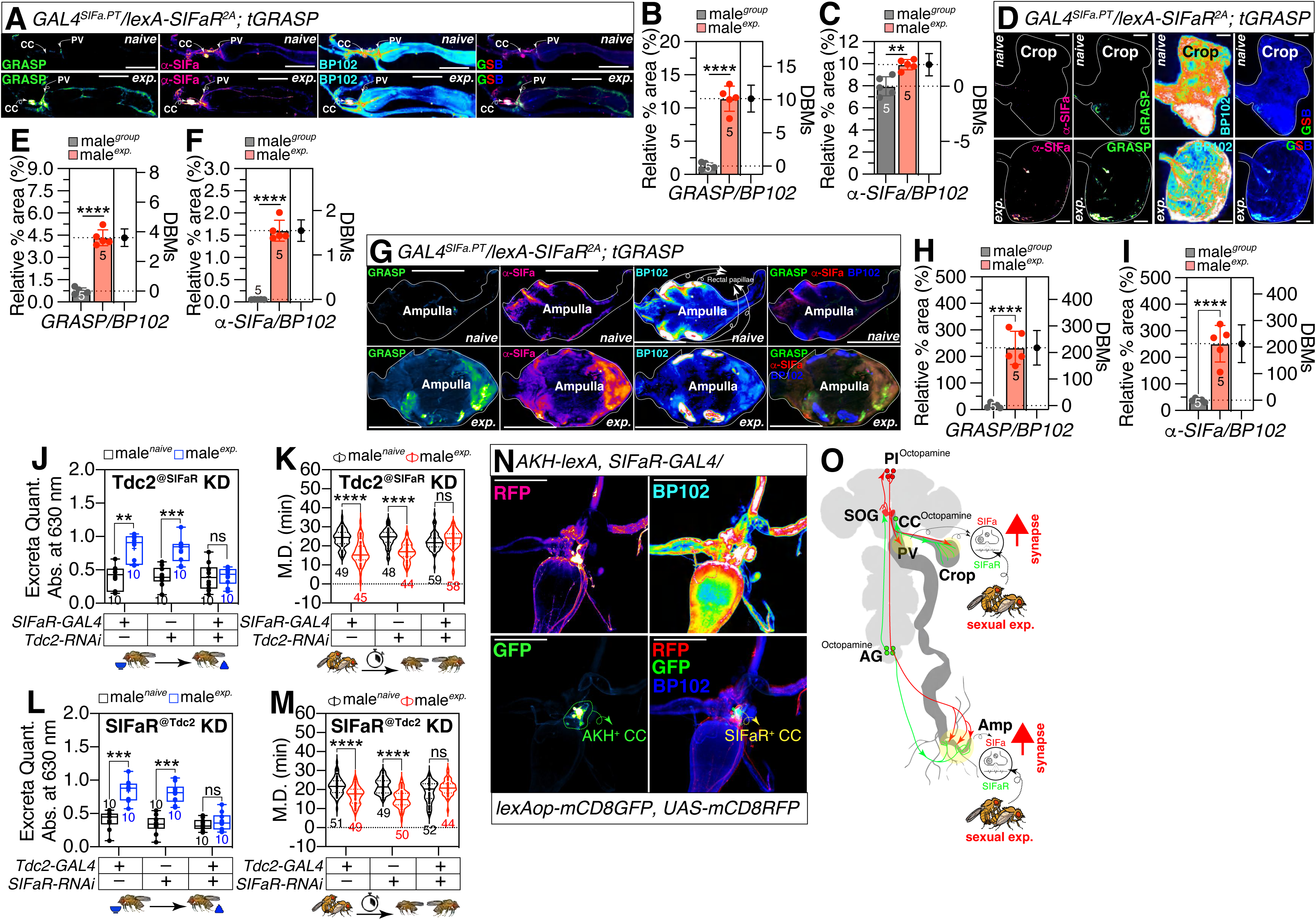
Mating enhances SIFa-SIFaR synaptic connectivity via an Octopaminergic/Tyraminergic mechanism in the Corpora Cardiaca. (A) tGRASP assay (resulted in a strong preferential GRASP signal in synaptic regions) for *GAL4^SIFa.PT^* and *lexA^SIFaR-2A^* in foregut of male flies in naïve (upper) and experienced (lower) conditions. SIFa peptide levels were measured using anti-SIFa antibody. Male flies expressing *GAL4^SIFa.PT^*, *lexA^SIFaR-2A^* and *lexAop-2-post-t-GRASP, UAS-pre-t-GRASP* were dissected after 5 days of growth. Synaptic transmission occurs from *GAL4^SIFa.PT^* to *lexA^SIFaR-2A^.* GFP is pseudo-colored as “green fire blue”. α-SIFa is pseudo-colored as “fire”. BP102 is pseudo-colored as “16 colors”. Scale bars represent 100 μm. (B-C) Quantification of relative value for synaptic intensity and SIFa peptide levels (two-tailed unpaired *t*-test). In all plots and statistical tests. Data are presented as mean ± s.e.m. ns = not significant (*p>0.05*), *\*p<0.05, **p<0.01, ***p< 0.001, ****p< 0.0001*. Sample sizes (n) are indicated in the figure panels. (D) tGRASP assay (resulted in a strong preferential GRASP signal in synaptic regions) for *GAL4^SIFa.PT^* and *lexA^SIFaR-2A^* in crop of male flies in naïve (upper) and experienced (lower) conditions. SIFa peptide levels were measured using anti-SIFa antibody. Male flies expressing *GAL4^SIFa.PT^*, *lexA^SIFaR-2A^*and *lexAop-2-post-t-GRASP, UAS-pre-t-GRASP* were dissected after 5 days of growth. Synaptic transmission occurs from *GAL4^SIFa.PT^* to *lexA^SIFaR-2A^.* GFP is pseudo-colored as “green fire blue”. α-SIFa is pseudo-colored as “fire”. BP102 is pseudo-colored as “16 colors”. Scale bars represent 100 μm. (E-F) Quantification of relative value for synaptic intensity and SIFa peptide levels (two-tailed unpaired *t*-test). In all plots and statistical tests. Data are presented as mean ± s.e.m. ns = not significant (*p>0.05*), *\*p<0.05, **p<0.01, ***p< 0.001, ****p< 0.0001*. Sample sizes (n) are indicated in the figure panels. (G) tGRASP assay (resulted in a strong preferential GRASP signal in synaptic regions) for *GAL4^SIFa.PT^* and *lexA^SIFaR-2A^* in ampulla of male flies in naïve (upper) and experienced (lower) conditions. SIFa peptide levels were measured using anti-SIFa antibody. Male flies expressing *GAL4^SIFa.PT^*, *lexA^SIFaR-2A^* and *lexAop-2-post-t-GRASP, UAS-pre-t-GRASP* were dissected after 5 days of growth. Synaptic transmission occurs from *GAL4^SIFa.PT^* to *lexA^SIFaR-2A^.* GFP is pseudo-colored as “green fire blue”. α-SIFa is pseudo-colored as “fire”. BP102 is pseudo-colored as “16 colors”. Scale bars represent 100 μm. (H-I) Quantification of relative value for synaptic intensity and SIFa peptide levels (two-tailed unpaired *t*-test). In all plots and statistical tests. Data are presented as mean ± s.e.m. ns = not significant (*p>0.05*), *\*p<0.05, **p<0.01, ***p< 0.001, ****p< 0.0001*. Sample sizes (n) are indicated in the figure panels. (J-M) Reciprocal genetic knockdown reveals functional coupling between SIFa signaling and Tdc2. Knockdown of Tdc2 in SIFaR-expressing neurons (J, K) or knockdown of SIFaR in Tdc2-expressing neurons (L, M) disrupts the mating-induced increase in excretion and abolishes the SMD phenotype, demonstrating that Tdc2 signaling is required for SIFa-dependent post-mating metabolic regulation. In all plots and statistical tests. Data are presented as mean ± s.e.m. ns = not significant *(p>0.05), *p<0.05, **p<0.01, ***p< 0.001, ****p< 0.0001*. Sample sizes (n) are indicated in the figure panels. (N) AKH and SIFaR show strong colocalization in CC. Male flies foregut expressing *SIFaR-GAL4* and *AKH-lexA* drivers together with *UAS-mCD8RFP* and *lexAop-mCD8GFP*. Scale bars represent 100 μm. (O) Schematic model of the SIFa-CC endocrine axis and behavioral modulation by SIFa-SIFaR synaptic plasticity. Mating enhances SIFa-SIFaR synapses in the CC, crop and ampulla. This activates two pathways: AKH to mobilize fat body energy, and OA/TA to regulate the crop and ampulla. Thus, synaptic plasticity at the CC, crop and ampulla orchestrates post-mating changes in both metabolism and gut function.

We next examined the neurochemical basis of this regulation. In insects, the octopaminergic/tyraminergic (OA/TA) system functions analogously to the mammalian noradrenergic system, acting as a key regulator of the “fight or flight” response, metabolic mobilization, and feeding homeostasis ^104^. We have previously reported that SIFa neurons modulate mating investment via Tdc2-dependent signaling, relying on Tyrosine decarboxylase 2 (Tdc2), the rate-limiting enzyme for OA/TA synthesis ^105^. To determine if this system operates within the SIFa-SIFaR vagus axis, we performed specific RNAi knockdown of *Tdc2* in SIFaR-expressing neurons. Strikingly, this depletion disrupted both the mating-induced increase in excretion and the SMD behavior (Fig. 4J–M).

Anatomical mapping provided a crucial distinction in how this chemical signal is deployed. The *Tdc2-GAL4* driver labels the crop and foregut regions heavily but is absent in the rectum (Fig. S4A). Since SIFaR neurons project to both regions, this suggests a functional subdivision: the OA/TA system largely functions within SIFaR-positive neurons originating from the CC (the anterior neurohemal organ), rather than the SIFaR neurons in AG. To confirm the identity of these anterior neurons, we used genetic intersection methods to verify that a subpopulation of CC neurons strongly expresses SIFaR (Fig. 4N). Notably, these SIFaR-positive CC neurons do not exhibit the overlapping brain projections seen in other interneurons (Fig. S4B), suggesting they primarily serve an endocrine role (Fig. 4O). We probed the interaction between SIFa and the primary metabolic hormone of the CC, Adipokinetic Hormone (AKH)— the insect homolog of glucagon. Knockdown of either AKH or SIFaR specifically within the CC revealed that AKH levels are tightly controlled by SIFa signaling via SIFaR (Fig. 5A–D). Whole-mount dissection of the CNS-gut axis showed that these AKH neurons extend dendrites restricted to the CC but project axons asymmetrically to the brain and VNC (Fig. 5E). Physiologically, mating triggered a drastic increase in both intracellular calcium (Ca^2+^) and AKH peptide levels within these cells (Fig. 5F– H), consistent with the mobilization of energy reserves required in the post-mating state.

**Figure. 5.**
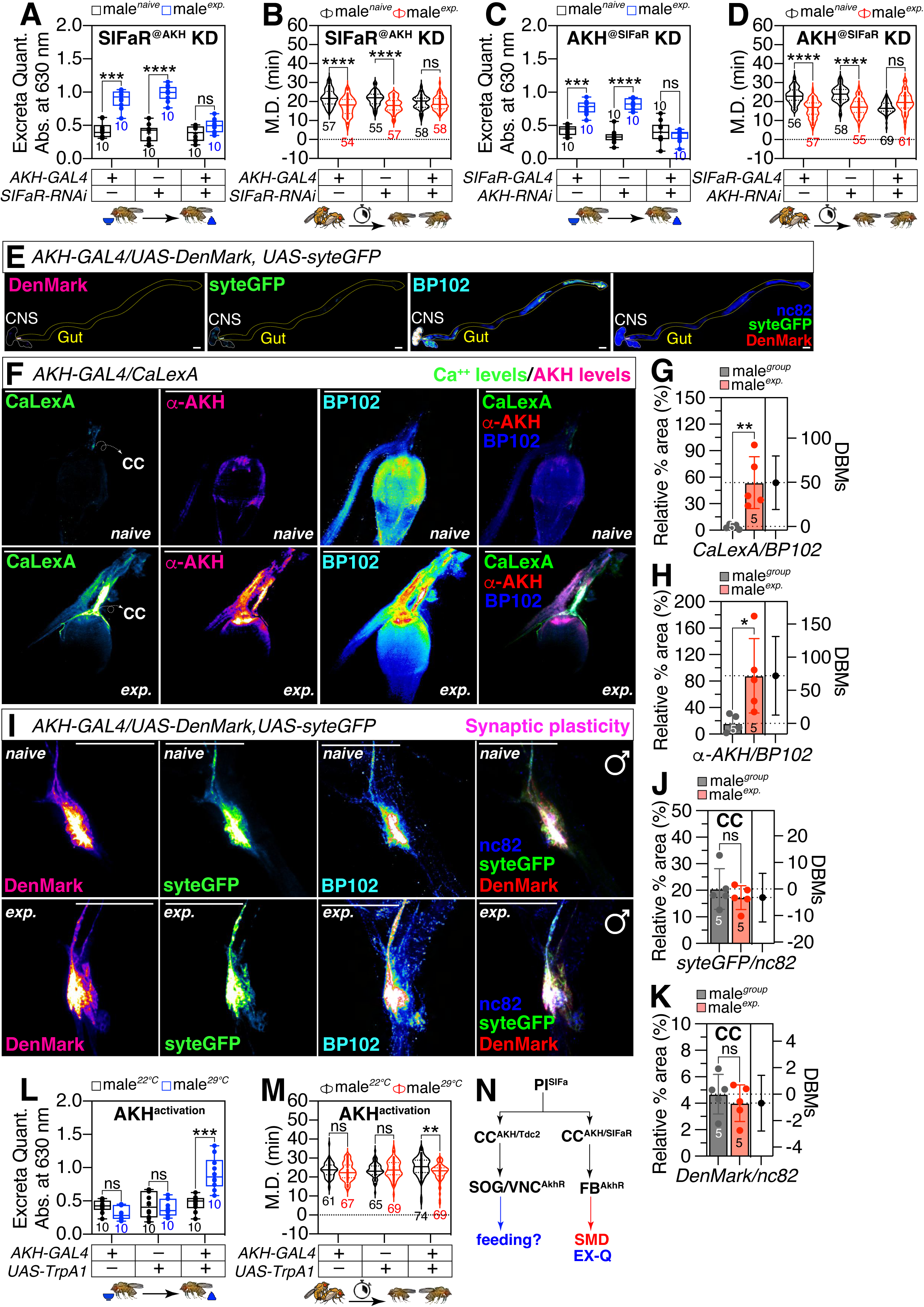
SIFa signaling regulates AKH secretion from the Corpora Cardiaca to drive post-mating metabolic homeostasis. (A-D) Reciprocal genetic knockdown reveals functional coupling between SIFa signaling and AKH. Knockdown of SIFaR in AKH expressing neurons (A, B) or knockdown of AKH in SIFaR-expressing neurons (C, D) disrupts the mating-induced increase in excretion and abolishes the SMD phenotype, demonstrating that AKH signaling is required for SIFa-dependent post-mating metabolic regulation. In all plots and statistical tests. Data are presented as mean ± s.e.m. ns = not significant *(p>0.05), *p<0.05, **p<0.01, ***p< 0.001, ****p< 0.0001*. Sample sizes (n) are indicated in the figure panels. (E) Male flies CNS and gut expressing *AKH-GAL4* together with *UAS-Denmark, UAS-syt.eGFP* were immunostained with anti-RFP (red) for dendrites, anti-GFP (green) for presynaptic terminals and anti-nc82 (blue) for neuropil. Scale bars represent 100 μm. (F) Different levels of neural activity of the brain as revealed by the CaLexA system in naïve, single and experienced flies. Male flies expressing *AKH-GAL4* along with *LexAop-CD2-GFP, UAS-mLexA-VP16-NFAT* and *LexAop-CD8-GFP-A2-CD8-GFP* were dissected after 5 days of growth (mated male flies had 1-day of sexual experience with virgin females). The dissected guts were then immunostained with anti-GFP (green), anti-AKH (red) and anti-BP102 (blue). GFP is pseudo-colored as “green fire blue”. α-AKH is pseudo-colored as “fire”. BP102 is pseudo-colored as “16 colors”. Scale bars represent 100 μm. (G-H) Quantification of relative intensity value for GFP fluorescence and AKH levels (two-tailed unpaired *t*-test). In all plots and statistical tests. Data are presented as mean ± s.e.m. ns = not significant (*p>0.05*), *\*p<0.05, **p<0.01, ***p< 0.001, ****p< 0.0001*. Sample sizes (n) are indicated in the figure panels. (I) Male flies ampulla expressing *AKH-GAL4* together with *UAS-Denmark* and *UAS-syt.eGFP* in naive (upper panels) and exp. (lower panels) conditions. Denmark is pseudo-colored as “fire”. GFP is pseudo-colored as “green fire blue”. BP102 is pseudo-colored as “16 colors”. Scale bars represent 100 μm. (J-K) Quantification of relative intensity value for Denmark and GFP fluorescence in CC (two-tailed unpaired *t*-test). In all plots and statistical tests. Data are presented as mean ± s.e.m. ns = not significant (*p>0.05*), *\*p<0.05, **p<0.01, ***p< 0.001, ****p< 0.0001*. Sample sizes (n) are indicated in the figure panels. (L-M) Genetic activation reveals the function of AKH. Activition of AKH induced the increase in excretion and induced the SMD phenotype. In all plots and statistical tests. Data are presented as mean ± s.e.m. ns = not significant *(p>0.05), *p<0.05, **p<0.01, ***p< 0.001, ****p< 0.0001*. Sample sizes (n) are indicated in the figure panels. (N) Schematic model of the SIFa-CC endocrine axis from CC to SOG/VNC and FB, modulating different behaviors.

Interestingly, while Ca^2+^ and AKH levels surged, AKH neurons did not exhibit the dramatic synaptic structural plasticity observed in the AG-to-RPs circuit (Fig. 5I–K). This stability is consistent with their projection patterns (Fig. S5A–C) and implies that mating modulates the AKH axis primarily via a secretion pathway rather than synaptic remodeling. To test this, we knocked down the AKH receptor (AkhR) in specific tissues. We found that AkhR knockdown in the Fat Body—the central storage organ for lipids and glycogen—disrupted SMD behavior, whereas neuronal knockdown did not (Fig. S5D). This confirms a neuroendocrine loop: SIFa signaling stimulates the CC (via SIFaR) to secrete AKH into the hemolymph, which then activates AkhR in the Fat Body to modulate energy homeostasis.

To map the broader connectivity of this module, we employed trans-Tango (an anterograde tracing tool that labels postsynaptic partners) and *retro*-Tango (a retrograde tool for labeling presynaptic inputs) ^106,107^. Trans-Tango analysis revealed that AKH neurons connect to downstream neurons projecting to the crop and ampulla, resembling the peripheral SIFaR projection pattern (Fig. S5E). Conversely, *retro*-Tango identified strong presynaptic inputs to AKH-CC neurons originating from the fan-shaped body and mushroom body (Fig. S5F–H). These regions, especially fan-shaped body are known targets of SIFa projections ^34,39,108,109^, strongly supporting a direct circuit where SIFa neurons integrate higher-order processing from the central complex to regulate the CC.

Finally, to demonstrate sufficiency, we artificially activated AKH neurons using dTrpA1, a temperature-sensitive cation channel that depolarizes neurons upon heating ^110–112^. Thermal activation was sufficient to mimic the mated state, inducing both increased excretion and reduced mating duration (Fig. 5L–M). We further confirmed that AKH neurons co-express Tdc2 and SIFaR (Fig. S5K) and utilize the OA/TA system to modulate these behaviors (Fig. S5I–J). Collectively, these data define a unified but bifurcated control system for the insect vagus homolog. SIFa neurons in the PI act as the central command. They project to the CC, activating SIFaR-positive neuroendocrine cells. These cells then execute a dual-modality program: (1) they utilize the OA/TA neurotransmitter system to drive downstream neurons regulating excreta and feeding homeostasis, and (2) they secrete AKH to signal the Fat Body, mobilizing energy to support the behavioral shift toward reduced mating investment (Fig. 5N). This architecture elegantly couples the immediate regulation of visceral physiology (excretion) with the systemic metabolic signaling (AKH/Fat Body) required to sustain the post-mating life-history transition.

### SIFa Signaling Orchestrates AstA-Mediated Metabolic Homeostasis via the Abdominal Ganglion

Feeding behavior is fundamental to energy homeostasis, a process tightly regulated by the neuropeptide Allatostatin A (AstA). AstA is a well-characterized satiety factor that inhibits feeding and regulates food choice based on nutritional value and metabolic state ^17,18,113^. Given the interplay between reproductive investment and metabolic demand, we hypothesized that the SIFa “vagus” network might recruit AstA signaling to coordinate energy balance ^114^. Confirming this interaction, reciprocal knockdown of AstA in SIFaR-expressing neurons, or SIFaR in AstA-expressing neurons, significantly disrupted the mating-induced increase in excretion and the SMD phenotype (Fig. 6A-D). Anatomical analysis revealed a strong colocalization of AstA and SIFaR in AG (Fig. 6E). These AG AstA/SIFaR neurons project extensive axons and dendrites to the RPs, paralleling SIFaR projections (Fig. 6F-H). This suggests that AG AstA neurons expressing SIFaR serve as a direct downstream target of descending SIFa signaling.

**Figure. 6.**
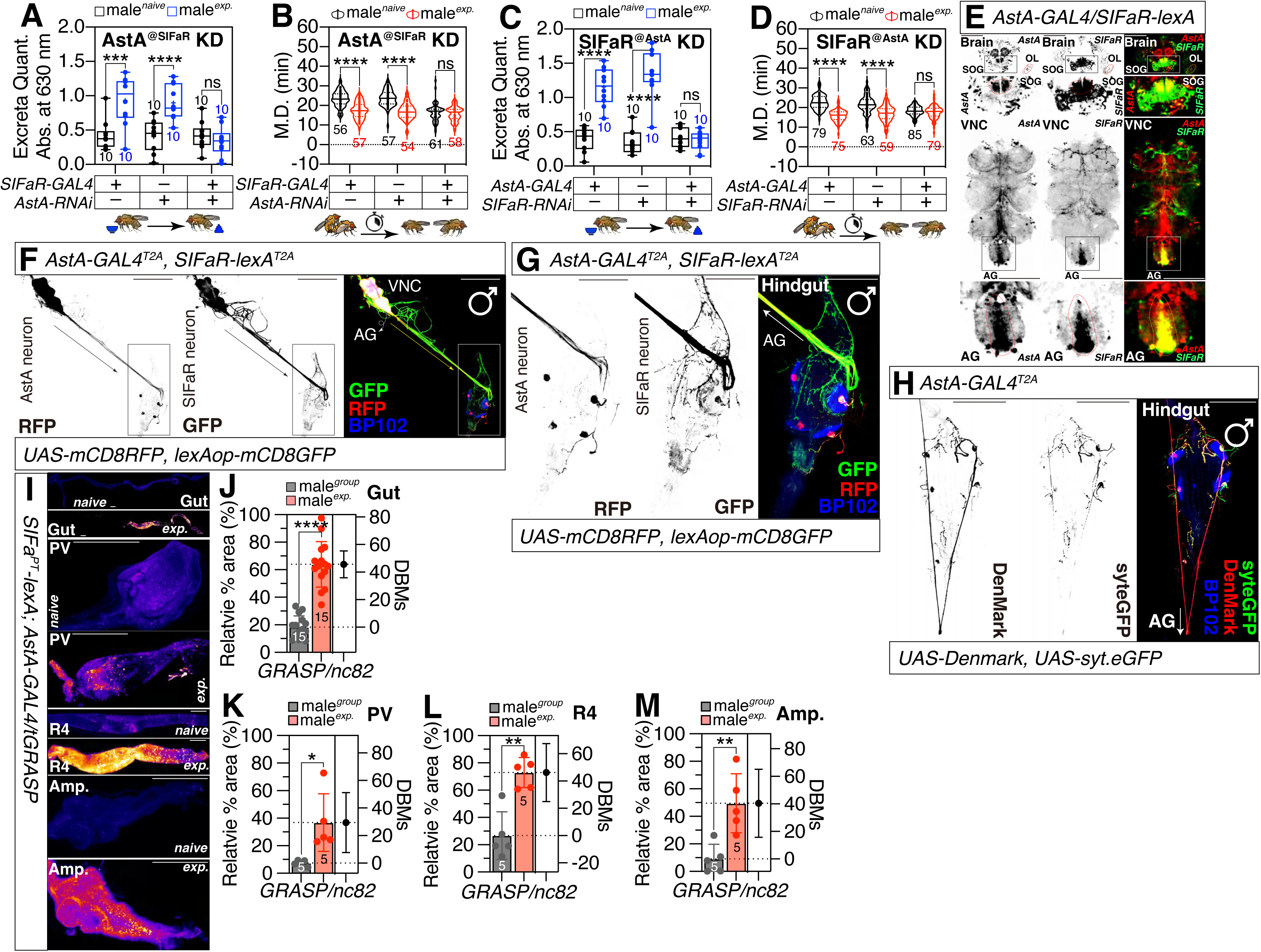
AstA neurons function downstream of SIFa signaling to couple feeding, metabolism, and reproductive state. (A-D) Reciprocal genetic knockdown reveals functional coupling between SIFa signaling and AstA. Knockdown of AstA in SIFaR-expressing neurons (A, B) or knockdown of SIFaR in AstA-expressing neurons (C, D) disrupts the mating-induced increase in excretion and abolishes the SMD phenotype, demonstrating that AstA signaling is required for SIFa-dependent post-mating metabolic regulation. In all plots and statistical tests. Data are presented as mean ± s.e.m. ns = not significant *(p>0.05), *p<0.05, **p<0.01, ***p< 0.001, ****p< 0.0001.* Sample sizes (n) are indicated in the figure panels. (E) AstA and SIFaR show strong colocalization in neurons of the Abdominal Ganglion (AG), identifying AG AstA/SIFaR neurons as a major site of signal integration between feeding and reproductive state. Brain, VNC and AG of male flies were immunostained with anti-RFP (red) for AstA and anti-GFP (green) for SIFaR. Scale bars represent 100 μm. (F-H) AG AstA/SIFaR neurons extend extensive axonal and dendritic projections to the reproductive processes, closely paralleling SIFaR projection patterns and supporting their role as direct downstream targets of descending SIFa signaling. VNC and hindgut of male flies expressing *AstA-GAL4^T2A^, SIFaR-lexA^T2A^* (F, G) together with *UAS-mCD8RFP* and *lexAop-mCD8GFP*. And *AstA-GAL4^T2A^*(H) together with *UAS-Denmark* and *UAS-syteGFP* were immunostained with anti-RFP (red), anti-GFP (green) and anti-BP102 (blue). Scale bars represent 100 μm. (I) Mating induces robust synaptic remodeling between SIFa and AstA neurons. tGRASP assay for *SIFa-lexA^PT^* and *AstA-GAL4* in whole region (upper panels), PV (second panels), R4 region (third panels) and ampulla regions of naïve (upper) and experienced (lower) male flies. Male flies expressing were dissected after 5 days of growth. Scale bars represent 100 μm. (J-M) Quantification of relative value for synaptic intensity between naïve and experienced male flies. The intensity of GFP fluorescence was normalized to that of the nc82. tGRASP analysis reveals a marked increase in synaptic connectivity in the foregut and hindgut regions following mating, indicating state-dependent strengthening of SIFa-AstA communication along the gut axis. In all plots and statistical tests. Data are presented as mean ± s.e.m. ns = not significant *(p>0.05), *p<0.05, **p<0.01, ***p< 0.001, ****p< 0.0001.* Sample sizes (n) are indicated in the figure panels.

While *Drosophila* enteroendocrine cells (EECs) in the R4 gut region express AstA ^19,115^, our projection mapping indicates that the AstA innervation of the hindgut originates not from these EECs, but from the AG AstA/SIFaR neurons (Fig. S6A-B). Furthermore, we identified previously uncharacterized projections from AG AstA neurons to the male reproductive system (Fig. S7A), implicating AstA in reproductive physiology beyond its established role in feeding. Notably, synaptic connectivity between SIFa and AstA neurons—visualized via tGRASP—dramatically increases in the foregut and hindgut regions following mating (Fig. 6I-M; Fig. S7B-E). This synaptic remodeling indicates that AstA-mediated feeding and metabolic regulation is actively orchestrated by the SIFa vagus network in a context-dependent manner.

### State-Dependent Synaptic Plasticity in the Fan-Shaped Body Reinforces the ‘Rest and Digest’ State

We next investigated how visceral information returns to the brain. AG-resident SIFaR/AstA neurons extend strong axonal projections ascending toward the Prow (PRW) region, located near the superior, lateral margin of the subesophageal zone (SEZ) (Fig. 7A). Using tGRASP to map SIFaR-to-SIFa connectivity, we detected strong synaptic contacts in the PRW, Pars Intercerebralis (PI), and the Fan-Shaped Body (FSB) (Fig. 7B; Fig. S8A-B). Interestingly, mating induced a spatial shift in this connectivity: ascending gut-to-brain synapses between SIFaR and SIFa neurons decreased in the PRW but dramatically increased in the FSB (Fig. 7C-D).

**Figure. 7.**
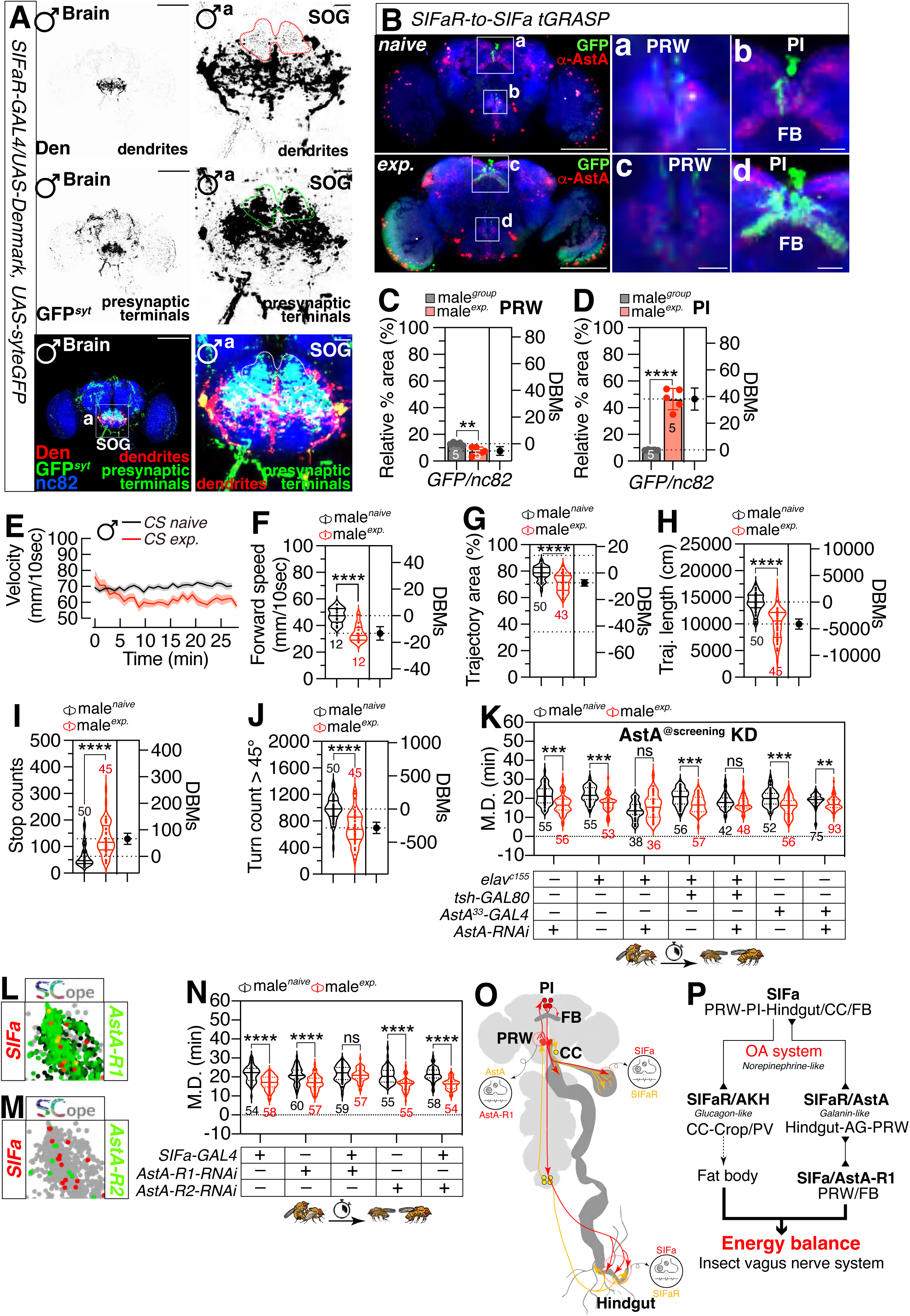
State-dependent synaptic plasticity redirects visceral feedback to the fan-shaped body to reinforce a ‘rest and digest’ state. (A) Male flies brain expressing *SIFaR-GAL4* together with *UAS-Denmark, UAS-syt.eGFP* were immunostained with anti-RFP (red) for dendrites, anti-GFP (green) for presynaptic terminals and anti-nc82 (blue) for neuropil. Scale bars represent 100 μm in brain panels and 20 μm in SOG panels. (B) tGRASP mapping reveals synaptic connectivity between SIFaR-expressing neurons and SIFa neurons in the PRW, PI, and FB regions. Mating induces a spatial redistribution of this connectivity, with decreased synaptic signal in the PRW and a pronounced increase in the FB. Scale bars represent 100 μm in brain panels and 20 μm in other panels. (C, D) Quantification of relative value for synaptic intensity between grouped and experienced male flies. The intensity of GFP fluorescence was normalized to that of the nc82. Quantification of tGRASP signal confirms mating-induced weakening of SIFaR-SIFa contacts in the PRW and strengthening in the FSB, indicating state-dependent synaptic remodeling along the gut-to-brain axis. In all plots and statistical tests. Data are presented as mean ± s.e.m. ns = not significant *(p>0.05), *p<0.05, **p<0.01, ***p< 0.001, ****p< 0.0001.* Sample sizes (n) are indicated in the figure panels. (E-J) Sexually experienced males display a hypo-active locomotor profile, characterized by reduced velocity (E, F), decreased trajectory area (G), shorter path length (H), increased stopping frequency (I), and reduced turning frequency (J), consistent with the engagement of FB-mediated circuits that promote energy conservation and a ‘rest and digest’ state. In all plots and statistical tests. Data are presented as mean ± s.e.m. ns = not significant *(p>0.05), *p<0.05, **p<0.01, ***p< 0.001, ****p< 0.0001.* Sample sizes (n) are indicated in the figure panels. For detailed methods, see the **MATERIALS AND METHODS** for a detailed description of the locomotion assays used in this study. (K) Mating duration of male flies for AstA knockdown screening. Disruption of AstA signaling in central neurons abolishes mating-induced changes in mating duration, identifying AstA as a critical mediator linking visceral state to behavioral output. In all plots and statistical tests. Data are presented as mean ± s.e.m. ns = not significant *(p>0.05), *p<0.05, **p<0.01, ***p< 0.001, ****p< 0.0001.* Sample sizes (n) are indicated in the figure panels. (L-M) Each tSNE visualization depicts the coexpression patterns of genes, with each color corresponding to the genes listed on the left, right, and bottom of the plot. The tissue name, as referenced on the Fly SCope website is indicated in the upper left corner of the tSNE plot. (N) Mating duration of male flies for AstA-R1 and R2 knockdown screening. In all plots and statistical tests. Data are presented as mean ± s.e.m. ns = not significant *(p>0.05), *p<0.05, **p<0.01, ***p< 0.001, ****p< 0.0001*. Sample sizes (n) are indicated in the figure panels. (O-P) Working model summarizing state-dependent routing of SIFa-dependent visceral feedback.

We hypothesize that the ascending connections to the PRW maintain basal homeostatic feeding regulation. Situated within the dorsal SEZ—the primary hub for gustatory integration and feeding motor control—neural activity in this region is ideally positioned to modulate reflexive ingestion and food acceptance mechanics^116–118^. In contrast, the mating-induced recruitment of the FSB signals a fundamental state transition toward energy conservation and recovery. The FSB is a higher-order integrative center within the Central Complex, essential for computing locomotor vectors, guiding visual navigation, and regulating sleep-wake arousal ^119–125^. Notably, accumulating evidence has established SIFa signaling as a potent promoter of sleep and a key regulator of energy balance and metabolic homeostasis^32,37,48,109,126^. We propose that the potentiation of SIFa synapses in the FSB reinforces the homeostatic ‘rest and digest’ drive, suppressing excessive exploratory locomotion to favor energy preservation and efficient internal resource management. Consistent with this, males with high sexual experience exhibited a hypo-active locomotor profile, moving slower (Fig. 7E-F), covering a smaller area (Fig. 7G) over shorter distances (Fig. 7H), with more frequent stops (Fig. 7I) and reduced turning frequency (Fig. 7J). These kinematic changes mirror the functional role of the FSB in modulating arousal thresholds and motor drive, likely promoting a quiescent or metabolically efficient state ^123^. We propose that the mating-induced synaptic potentiation from SIFaR neurons to SIFa neurons in the FSB urges the male to consolidate the “rest and digest” state, thereby securing energy reserves for future mating opportunities ^127,128^.

### Modular Control of Excretion and Behavior by Neuronal versus Enteric AstA

AstA is expressed across the central nervous system (brain and VNC) and the gut ^9,16^. While brain and gut AstA functions are documented, the specific contribution of AG AstA remains unclear ^19,129^. To dissect these distinct pools, we utilized *tsh-GAL80* to inhibit GAL4 expression in the VNC, and the *AstA*^33^*-GAL4* driver to specifically label gut EECs. We found that both neuronal and gut-derived AstA are required for the mating-induced increase in excretion (Fig. S8C). However, the behavioral switch (SMD) strictly requires neuronal AstA, whereas gut EEC AstA is dispensable (Fig. 7K). This dissociation suggests a modular system where endocrine AstA regulates physiological waste clearance, while neuronal AstA integrates metabolic states to modulate reproductive behavioral investment.

### A Bidirectional Feedback Loop via AstA Receptors

AstA signaling is transduced by two receptors, AstA-R1 and AstA-R2, which exhibit distinct expression profiles. Using CCT-T2A-KI lines ^95^, we confirmed that *AstA-R1* is broadly expressed throughout the CNS, overlapping significantly with SIFaR. In contrast, *AstA-R2* expression is highly restricted to specific integrative centers, including the Optic Lobe (OL), PI, and AG (Fig. S7H-I).

To determine the central role of these receptors, we employed an intersectional genetic approach using *otd-FLP* combined with *tub-GAL80* flanked by FRT sites ^130,131^. In this system, *otd*-driven FLPase removes the GAL80 repressor specifically in the brain, allowing us to isolate brain-specific receptor function. We found that expression of both AstA-R1 and AstA-R2 in the brain is required to generate the post-mating excretory increase and the modulation of mating investment (Fig. S7F-G).

Finally, we explored the circuit architecture closing the loop between the gut and the SIFa network. Since AG SIFaR/AstA neurons project ascending axons to the PRW— a region rich in SIFa dendrites (Fig. 7A-B) ^36,37^—we hypothesized that AstA provides direct feedback to SIFa neurons. Single-cell transcriptomic data (SCope) ^96^ predicts that SIFa neurons express *AstA-R1* but not *AstA-R2* (Fig. 7L-M). Validating this connection, RNAi-mediated knockdown of *AstA-R1* specifically in SIFa neurons disrupted SMD behavior, whereas *AstA-R2* knockdown did not (Fig. 7N). This confirms a bidirectional circuit: SIFa descends to the gut to regulate AstA release, and AstA signals back to SIFa neurons via AstA-R1, reinforcing the logic of a vagus-like sensorimotor loop ^126^ (Fig. 7O).

### SIFa is an Evolutionarily Conserved “Insect Vagus Nerve”

Insects exhibit immense biodiversity, with the five major orders—Hemiptera (true bugs), Lepidoptera (butterflies/moths), Diptera (flies), Hymenoptera (wasps/ants), and Coleoptera (beetles)—encompassing over 100,000 described species each ^132^. To investigate whether the SIFa network represents a universally conserved “insect vagus nerve,” we analyzed SIFa morphology across the insect kingdom. We examined the Brown Planthopper (*Nilaparvata lugens, Hemiptera*), Fall Armyworm (*Spodoptera frugiperda, Lepidoptera*), Harvest Ant (*Messor structor, Hymenoptera*), Cricket (*Acheta domesticus, Orthoptera*), Honey bee (*Apis mellifera, Hymenoptera*) and Seven-spot Ladybug (*Coccinella septempunctata, Coleoptera*), alongside other *Dipteran* species (*D. virilis, D. yakuba, D. pseudoobscura, D. mojavensis*) (Fig. S9).

SIFa neuropeptide sequence and function are known to be highly conserved ^29^. Indeed, recent comparative molecular analyses have further demonstrated the profound sequence and structural preservation of the SIFamide signaling machinery across divergent insect orders^133^. To extend these molecular insights to the level of functional neuroanatomy, we analyzed SIFa morphology across the insect kingdom. Using our specific SIFa antibodies, we observed that every species tested displayed a striking conservation of neuroanatomy: distinct, well-branched brain SIFa neurons sending long, direct projections to the hindgut (Fig. S9). This structural ubiquity supports the model that SIFa neurons function as a universal ‘insect vagus nerve.’ We propose that this network orchestrates the essential ‘rest and digest’ state by recruiting the AstA (Cholecystokinin-like) system ^9,134^ to promote satiety and sedation, while actively balancing metabolic homeostasis against the catabolic drives of the Octopamine (Norepinephrine-like) ^135^ and AKH (Glucagon-like) systems ^100,136^ (Fig. 7P).

## DISCUSSION

In this study, we establish the SIFamide (SIFa) peptidergic circuit as the functional evolutionary homolog of the vertebrate vagus nerve, resolving a longstanding question regarding the neural substrate of the insect gut-brain axis. We demonstrate that SIFa neurons, originating in the PI, project the longest identified axons in the *Drosophila* nervous system to directly innervate the hindgut ampulla and the CC, bypassing the segmental logic of the ventral nerve cord. This direct brain-to-viscera connection forms a bidirectional loop with SIFaR and AstA neurons in the abdominal ganglion, which relay metabolic status back to the brain’s PWR and FSB regions to modulate behavioral state. We further show that this circuit is highly plastic; sexual satiety strengthens the brain-hindgut synaptic connection via an OA/TA gating mechanism, shifting the organism’s priority from reproductive investment (mating) to metabolic replenishment accompanied by enhanced excretory processing. By regulating AKH secretion and hindgut motility, the SIFa system integrates metabolic homeostasis with life-history trade-offs, a functional architecture we found conserved across *Hemiptera*, *Lepidoptera*, and *Hymenoptera*.

An intriguing aspect of the insect vagus nerve is its pronounced sexual dimorphism. We found that the SIFa-dependent potentiation of excretion and the shift toward metabolic homeostasis are entirely male-specific^60,137^. This asymmetry likely reflects the divergent reproductive economics of the sexes. For the male, successful copulation imposes a severe metabolic toll and depletes finite seminal resources, necessitating an immediate vagal-mediated switch to a ‘rest and digest’ state to ensure future mating viability^59,65^. In contrast, the female post-mating response is predominantly governed by the Sex Peptide (SP) pathway^63,138^, which rapidly remodeling feeding, digestion, and oviposition to prioritize egg production rather than somatic energy replenishment^74,139^. It appears that while females utilize SP-mediated endocrine cascades to process the incoming ejaculate, males employ the SIFa vagus-like circuit as an active coping mechanism for ejaculate expenditure. This dichotomy highlights the evolutionary flexibility of the gut-brain axis, demonstrating that even highly conserved neural substrates can be sexually repurposed to meet distinct life-history strategies.

Moreover, we uncover an unanticipated degree of plasticity within the SIFa system itself. Sexual experience not only strengthens the synaptic output from SIFa to SIFaR neurons but also robustly enhances the dendritic and axonal arbors of the central SIFa neurons at the hindgut ampulla. This suggests that the “command” neurons of the insect vagus actively remodel their own peripheral interface to match the animal’s reproductive history, a feature that may be critical for fine-tuning the rest-and-digest response. The identification of the SIFa system as the insect vagus nerve challenges the traditional view that the stomatogastric nervous system (SNS), specifically the recurrent nerve, is the sole autonomic regulator in insects ^140^. While the recurrent nerve manages local deglutition patterns, it lacks the systemic reach and central integration characteristic of the vertebrate vagus. A true vagal homolog must satisfy specific anatomical criteria: it must originate in a central neuroendocrine command center and physically connect the brain to the distal visceral periphery. SIFa neurons fulfill this requirement precisely. Unlike the recurrent nerve, which originates in the peripheral frontal ganglion and largely terminates at the anterior midgut, SIFa neurons originate in the PI—a region ontogenetically and functionally analogous to the vertebrate hypothalamus and the dorsal motor nucleus of the vagus ^141^. From this central command center, SIFa axons traverse the entire length of the ventral nerve cord to form a dense plexus on the hindgut ampulla, providing the “long leash” innervation required to coordinate defecation and osmoregulation with central brain states. This morphology represents a direct physical link between the brain’s decision-making centers and the metabolic periphery, functionally mirroring the vagus nerve’s projection from the medulla to the distal colon.

Beyond anatomy, the SIFa system recapitulates the physiological duality of the vertebrate autonomic nervous system, specifically the “rest and digest” functions of the parasympathetic division. As highlighted in recent comprehensive syntheses, the vertebrate vagus nerve serves as a central hub for promoting anabolic states and suppressing sympathetic arousal to maintain energy homeostasis^55^. The vertebrate vagus nerve promotes anabolic states by stimulating gastric motility and regulating pancreatic hormones while suppressing high-arousal sympathetic states ^45^. Similarly, we found that SIFa signaling drives the post-mating “rest and digest” phase by coordinating two distinct effector pathways. First, it modulates the release of AKH (the insect glucagon) from the CC, thereby controlling lipid mobilization and energy storage in the fat body (Fig. 5). This interaction with the CC and adjacent Insulin-Producing Cells (IPCs) mirrors the vagal regulation of insulin and glucagon secretion from the vertebrate pancreas ^142^. Second, SIFa acts as a potent regulator of behavioral state, promoting sleep and facilitating digestive/metabolic processing while actively suppressing reproductive behaviors such as mating duration ^36–38^. This mutual inhibition—where anabolic recovery drives suppress reproductive investment— defines the autonomic logic of the SIFa circuit, ensuring the animal does not engage in energetically expensive behaviors like mating when metabolic restoration is required.

Crucially, the vertebrate vagus nerve is a bidirectional superhighway, with afferent sensory fibers constituting the majority of its axons to convey visceral feedback to the brain. Our data reveal that the SIFa circuit possesses an equivalent sensory arm mediated by the SIFaR/AstA neurons in the abdominal ganglion. We demonstrate that these peripheral neurons, which innervate the hindgut, project ascending axons back to the Subesophageal Zone (SOG/PWR) and the FSB of the central brain. This feedback loop allows the system to integrate “interoceptive” cues regarding gut distension and nutritional status directly with gustatory processing and locomotor control. Specifically, the interaction between SIFa and AstA---a peptide analogous to vertebrate satiety signals like CCK---creates a push-pull regulatory system that fine-tunes metabolic homeostasis ^134^. SIFa signaling promotes the ‘rest and digest’ state, facilitating energy conservation and metabolic recovery, while AstA feedback reinforces satiety and modulates feeding cessation according to internal energy status. This reciprocal circuitry mirrors the integration of gastric distension and nutrient sensing in the vertebrate Nucleus of the Solitary Tract (NTS), enabling the organism to fine-tune its behavior based on real-time physiological feedback ^143^.

Recent studies have highlighted the foundational role of specific neural populations in coordinating global brain states, further supporting the central integrative function of the SIFa circuit. Previous work demonstrated that a discrete population of neurons expressing the cation channel Trpg templates patterned, stimulus-independent neural activity (PSINA) across the entire *Drosophila* brain. Building upon this circuit architecture ^144^, recent findings have revealed that four SIFamide (SIFa)-expressing neurons actively pattern this brain-wide PSINA template through SIFa neuropeptide signaling ^41^. Together, these studies establish that SIFa neurons function as master orchestrators of brain neuronal internal states. However, a notable limitation of this prior work is that the investigations into the SIFa and Trpg networks were strictly restricted to the pupal developmental stage. Our current findings address this limitation by extending the functional relevance of the SIFa network into adulthood, demonstrating that these central command neurons continue to operate as a vital, highly plastic “insect vagus nerve” to coordinate visceral physiology and manage metabolic homeostasis throughout the mature organism’s lifespan.

Finally, the evolutionary conservation of this neuroanatomical framework underscores its fundamental importance in insect biology. The presence of SIFa hindgut projections in diverse species, including *Rhodnius prolixus* (Hemiptera), *Chilo suppressalis* (Lepidoptera), and *Messor structor* (Hymenoptera), suggests that the “Insect Vagus” is not a dipteran adaptation but an ancestral feature of the class Insecta. In social insects like ants, this axis likely expanded to regulate colony-level metabolic needs, such as trophallaxis, effectively functioning as a “social vagus” ^145–150^. The SIFa system thus represents a unified solution to the universal biological problem of energy homeostasis: a centralized, bidirectional interface that couples the internal metabolic reality of the gut with the external behavioral drives of the brain ^3,4,11,16,151–154^.

## MATERIALS AND METHODS

### Fly Stocks and Husbandry

*Drosophila melanogaster* were raised on cornmeal-yeast medium at similar densities to yield adults with similar body sizes. Flies were kept in 12 h light: 12 h dark cycles (LD) at 25□ (ZT 0 is the beginning of the light phase, ZT12 beginning of the dark phase) except for some experimental manipulation. Wild-type flies were *Canton-S* (*CS*).

Following lines used in this study, *Canton-S* (#64349), *Oregon-R-C (#5), Df(1)Exel6234* (#7708), *SIFa-RNAi (#60484)*, *UAS-tdTomato* (#32222), *elav^c155^; UAS-Dicer* (#25750), *tsh-GAL80* (#605556), *lexAop-tdTomato.nls* (#66690), *lexAop-mCD8GFP* (#32205), *lexAop-CD8GFP; UAS-mLexA-VP16-NFAT, lexAop-rCD2-GFP* (#66542), *UAS-post-t-GRASP, lexAop2-pre-t-GRASP* (#79039), *UAS-pre-t-GRASP, lexAop2-post-t-GRASP* (#79040), *GAL4^24F06^* (#49087), *SIFaR-RNAi^JF01849^*(#25831), *SIFaR-RNAi^HMS00299^* (#34947), *lexA^24F06^* (#52695), *Tdc2-GAL4* (#9313), *Tdc2-RNAi* (#25871), *AKH-RNAi* (#27031), *Lsp^2.1^-GAL4* (#6357) *UAS-TrpA1* (#61504), AstA-RNAi (#25866), *AstA-R1-RNAi* (#27280), *AstA-R2-RNAi* (#25935), *AstA-R2-GAL4* (#84594), *AstA-R1-GAL4* (#84709), *tub-GAL80^ts^* (#7108), *UAS-CD4tdGFP* (#35839), *UAS-RedStinger* (#8546), *UAS-Denmark, UAS-syt.eGFP* (#33065), *UAS-KCNJ2* (#6596), *UAS-mCD8RFP, lexAop-mCD8GFP* (#32229), *elav^c155^; Otd^FLP^/CyO; tub-FRT-GAL80* (#38881), *UAS-mCD8GFP, QUAS-mtdTomato; trans-tango* (#95317), and *QUAS-mtdTomato; retro-tango; UAS-retro-tango* (#99661) were obtained from the Bloomington *Drosophila* Stock Center at Indiana University. The following lines, *AKH-lexA* (#FBF00134), *AstA-GAL4* (#FBA00006), *SIFaR-GAL4^T2A^* (#FBF00102), and *SIFaR-lexA^T2A^* (#FBF00086) were obtained from Qidong Fungene Biotechnology in China. The *GAL4^SIFa.PT^* was a gift from Dr. Jan A. Veenstra. The *w^CS^* was a gift from Dr. Hongtao Qin (Hunan University). The *AstA^33^-GAL4* was a gift from Drs. Wei Song (Wuhan University). *The SIFa^PT^-lexA* has been described elsewhere^36–38^.

The *CS* background was selected as the experimental background due to its well-characterized and consistent SMD behaviors. To ensure that genetic variation did not confound our results, all GAL4, UAS, and RNAi lines employed in our assays were rigorously backcrossed into the *CS* strain, often exceeding ten generations of backcrossing. This approach was undertaken to isolate the effects of our genetic manipulations from those of genetic background. We assert that the extensive backcrossing to the *CS* background, in concert with the internal control in SMD, provides a stable platform for the accurate interpretation of the SMD phenotypes observed in our experiments. To reduce the variation from genetic background, all flies were backcrossed for at least 10 generations to *CS* strain. For the generation of outcrosses, all GAL4, UAS, and RNAi lines employed as the virgin female stock were backcrossed to the *CS* genetic background for a minimum of ten generations. Notably, the majority of these lines have been maintained in a *CS* backcrossed state for long-term generations subsequent to the initial outcrossing process, exceeding ten backcrosses. Based on our experimental observations, the genetic background of primary significance is that of the X chromosome inherited from the female parent. Consequently, we consistently utilized these fully outcrossed females as virgins for the execution of experiments pertaining to SMD behavior. We have previously demonstrated that the genetic background exerts negligible influence on SMD behavior, as reported in our prior publication ^155^. The mutants and transgenic lines utilized in this study have been previously characterized, with the exception of the novel transgenic strains that we generated and describe herein.

### Mating duration assay

The mating duration assay in this study has been reported ^155–159^. To enhance the efficiency of the mating duration assay, we utilized the *Df(1)Exel6234* (DF here after) genetic modified fly line in this study, which harbors a deletion of a specific genomic region that includes the sex peptide receptor (SPR) ^138,160^. Previous studies have demonstrated that virgin females of this line exhibit increased receptivity to males ^138^. We conducted a comparative analysis between the virgin females of this line and the *CS* virgin females and found that both groups induced SMD. Consequently, we have elected to employ virgin females from this modified line in all subsequent studies. For group reared (naïve) males, 40 males from the same strain were placed into a vial with food for 5 days. For experienced males, 40 males from the same strain were placed into a vial with food for 4 days then 80 DF virgin females were introduced into vials for last 1 day before assay. 40 DF virgin females were collected from bottles and placed into a vial for 5 days. These females provide both sexually experienced partners and mating partners for mating duration assays. At the fifth day after eclosion, males of the appropriate strain and DF virgin females were mildly anaesthetized by CO_2_. After placing a single female in to the mating chamber, we inserted a transparent film then placed a single male to the other side of the film in each chamber. After allowing for 1 h of recovery in the mating chamber in 25□ incubators, we removed the transparent film and recorded the mating activities. Only those males that succeeded to mate within 1 h were included for analyses. Initiation and completion of copulation were recorded with an accuracy of 10 sec, and total mating duration was calculated for each couple. Part of genetic controls with *GAL4/+* or *UAS/+* lines were omitted from supplementary figures, as prior data confirm their consistent exhibition of normal SMD behavior ^155–157,161,162^. Hence, genetic controls for SMD behavior were incorporated exclusively when assessing novel fly strains that had not previously been examined. In essence, internal controls were predominantly employed in the experiments, as SMD behavior exhibit enhanced statistical significance when internally controlled. In SMD experiments, the naïve condition and sexually experienced males act as mutual internal controls for one another. A statistically significant divergence between naïve and experienced males indicates that the experimental procedure does not alter SMD. Conversely, the absence of a statistically significant difference suggests that the manipulation does impact SMD. All assays were performed from noon to 4 PM. We conducted blinded studies for every test.

### Antibodies and reagents

To generate the SIFa-antibody and AstA-antibody, the peptide sequence AYRKPPFNGSIFa of SIFa^37^ and SRPYSFGLa of AstA was synthesized and conjugated to Keyhole Limpet Hemocyanin (KLH). The efficiency of the conjugation was confirmed by SDS-PAGE analysis. Healthy rabbits were immunized via multiple subcutaneous injections with the peptide-KLH conjugate mixed with Freund’s adjuvant. The immunization schedule included a primary immunization and two booster injections, each separated by three weeks. Two weeks after the final booster, serum was collected from the rabbits via cardiac puncture. The serum was purified using affinity chromatography, and the antibody titer and specificity were determined by ELISA. Antibody synthesis was performed by Wuhan Vapol Bioscience Company (http://www.vapolbio.com).

### Immunostaining

The dissection and immunostaining protocols for the experiments are described elsewhere ^157^. After 5 days of eclosion, the *Drosophila* brain was taken from adult flies and fixed in 4% formaldehyde at room temperature for 30 minutes. The sample was washed three times (5 minutes each) in 1% PBT and then blocked in 5% normal goat serum for 30 minutes. The sample next be incubated overnight at 4□ with primary antibodies in 1% PBT, followed by the addition of fluorophore-conjugated secondary antibodies for one hour at room temperature. The brain was mounted on plates with an antifade mounting solution (Solarbio) for imaging purposes.

For the brain and hindgut of other insect species, the following protocol was applied. Dissection was performed in phosphate-buffered saline (PBS). Tissues were fixed in 4% formaldehyde for 30 minutes at room temperature; for thicker tissues such as the brain of *Spodoptera frugiperda*, fixation time was extended to 1 hour. After fixation, samples were washed four times (≥10 minutes each) in PBT. Subsequently, tissues were blocked in 5% normal goat serum for 30 minutes at room temperature. Primary antibodies diluted in PBT were applied overnight at 4°C. Following the same washing steps (four times in PBT, ≥10 minutes each), samples were incubated with fluorophore-conjugated secondary antibodies overnight at 4°C. After a final washing series (four times in PBT, ≥10 minutes each), tissues were mounted on slides with an antifade mounting solution (Solarbio) for imaging.

Samples were imaged with Zeiss LSM880. Antibodies were used at the following dilutions: Chicken anti-GFP (1:500, Invitrogen), mouse anti-nc82 (1:50, DSHB), rabbit anti-RFP (1:500, Rockland Immunochemicals), rabbit anti-AKH (1:200, Kerafast), mouse anti-BP102 (1:200, DSHB), mouse anti-nc82 (1:50, DSHB), rabbit anti-SIFa (generated by this study) ^37^ (1:200), anti-AstA (1:200, generated by this study), Alexa-488 donkey anti-chicken (1:200, Jackson ImmunoResearch), Alexa-555 goat anti-rabbit (1:200, Invitrogen), Alexa-647 goat anti-mouse (1:200, Jackson ImmunoResearch). For F-actin staining, samples were incubated with Alexa Fluor™ 647-conjugated phalloidin (Invitrogen) at a dilution of 1:200 for 1 hour at room temperature in the dark.

### Quantitative analysis of fluorescence intensity

To ascertain calcium levels and synaptic intensity from microscopic images, we dissected and imaged five-day-old flies of various social conditions and genotypes under uniform conditions. For group reared (naïve) flies, the flies were reared in group condition and dissect right after 5 days of rearing without any further action. For sexual experienced flies, the flies were reared in group condition after 4 days of rearing and will be given virgins to give them sexual experience for one day, those flies will also be dissected at the same time as group and single reared flies after one day. The GFP signal was amplified through immunostaining with chicken anti-GFP primary antibody. Image analysis was conducted using ImageJ software. For the quantification of fluorescence intensities, an investigator, blinded to the fly’s genotype, thresholded the sum of all pixel intensities within a sub-stack to optimize the signal- to-noise ratio, following established methods ^163^. The total fluorescent area or region of interest (ROI) was then quantified using ImageJ, as previously reported. For CaLexA signal quantification, we adhered to protocols detailed by Kayser et al. ^164^, which involve measuring the ROI’s GFP-labeled area by summing pixel values across the image stack. This method assumes that changes in the GFP-labeled area are indicative of alterations in the CaLexA signal, reflecting synaptic activity. ROI intensities were background-corrected by measuring and subtracting the fluorescent intensity from a non-specific adjacent area, as per Kayser et al. ^164^. For the analysis of t-GRASP signals, which specifically reconstitute fluorescence only at synaptic interfaces rather than within the cytoplasm^163^, a 3D sub-stack encompassing all discrete synaptic puncta was extracted. To ensure that only bona fide synaptic contacts were quantified, a uniform threshold was applied by a genotype-blinded investigator to rigorously eliminate any residual non-synaptic background fluorescence and achieve an optimal signal-to-noise ratio. The total fluorescence area within each ROI—which directly corresponds to the area of synaptic contact—was quantified using ImageJ. While the computational workflow for thresholding and area measurement employed a similar macro-based approach to that used for CaLexA quantification, the t-GRASP readout specifically reports structural synaptic occupancy rather than transcriptional activity.

### Particle analysis

Before the particle analysis, an investigator, blinded to the fly’s genotype, thresholded the sum of all pixel intensities within a sub-stack to optimize the signal-to-noise ratio, following established methods ^163^. To quantitatively measure particle intensity of cell number and synaptic puncta in microscopic images, we applied ImageJ software. Initially, the image is converted to grayscale to reduce complexity and enhance contrast. Subsequently, thresholding techniques are employed to binarize the image, distinguishing particles from the background. This binarization can be achieved through automated thresholding algorithms or manual adjustment to optimize the segmentation. The results of these measurements are then available for review by conducting “Analyze Particles” function of ImageJ. All specimens were imaged under identical conditions.

### Single-nucleus RNA-sequencing analyses

The snRNAseq dataset analyzed in this paper is published in ^165^ and available at the Nextflow pipelines (VSN, https://github.com/vib-singlecell-nf), the availability of raw and processed datasets for users to explore, and the development of a crowd-annotation platform with voting, comments, and references through SCope (https://flycellatlas.org/scope), linked to an online analysis platform in ASAP (https://asap.epfl.ch/fca). For the generation of the tSNE plots, we utilized the Fly SCope website (https://scope.aertslab.org/#/FlyCellAtlas/*/welcome). Within the session interface, we selected the appropriate tissues and configured the parameters as follows: ‘Log transform’ enabled, ‘CPM normalize’ enabled, ‘Expression-based plotting’ enabled, ‘Show labels’ enabled, ‘Dissociate viewers’ enabled, and both ‘Point size’ and ‘Point alpha level’ set to maximum. For all tissues, we referred to the individual tissue sessions within the ‘10X Cross-tissue’ RNAseq dataset. Each tSNE visualization depicts the coexpression patterns of genes, with each color corresponding to the genes listed on the left, right, and bottom of the plot. The tissue name, as referenced on the Fly SCope website is indicated in the upper left corner of the tSNE plot. Dashed lines denote the significant overlap of cell populations annotated by the respective genes. Coexpression between genes or annotated tissues is visually represented by differentially colored cell populations. For instance, yellow cells indicate the coexpression of a gene (or annotated tissue) with red color and another gene (or annotated tissue) with green color. Cyan cells signify coexpression between green and blue, purple cells for red and blue, and white cells for the coexpression of all three colors (red, green, and blue). Consistency in the tSNE plot visualization is preserved across all figures.

Single-cell RNA sequencing (scRNA-seq) data from the *Drosophila melanogaster* were obtained from the Fly Cell Atlas website (https://doi.org/10.1126/science.abk2432). Oenocytes gene expression analysis employed UMI (Unique Molecular Identifier) data extracted from the 10x VSN oenocyte (Stringent) loom and h5ad file, encompassing a total of 506,660 cells. The Seurat (v4.2.2) package (https://doi.org/10.1016/j.cell.2021.04.048) was utilized for data analysis. Violin plots were generated using the “Vlnplot” function, the cell types are split by FCA.

### Ex-Q assay

The EX-Q assays were conducted according to previously established protocols ^77^. In short, 20 flies were placed in a tube containing a food mixture composed of 5% sucrose, 5% yeast, and 1% agar, supplemented with 1% erioglaucine disodium (blue dye). Following a 48-hour adaptation period to the blue-dyed food, the flies were transferred to individual tubes containing the same dyed food. They were allowed to feed and excrete for 24 hours, after which the food was removed for 3 hours to clear any remaining gut contents. The flies were then removed, and the dye was dissolved in 2 mL of water. The absorbance of the resulting solution was measured at 630 nm^166^ For TrpA1 and shi^ts^ thermogenetic experiments, flies were maintained in 22°C and transferred to 29°C during dye food feeding.

### Locomotion assay

To detect and quantify the activity of flies, we have developed the Fly Trajectory Dynamics Tracking (FlyTrDT) software ^167^. This is an open-source, custom-written Python program that utilizes the free OpenCV machine vision library and the Python Qt library. The FlyTrDT software simultaneously records the trajectory information of each fly and calculates various indicators of the group over a certain period. For each frame acquired, the moving fly is segmented using the binarization function from the OpenCV library. Subsequently, a Gaussian blur and morphological closing and opening operations were performed on the extracted foreground pixels to consolidate detected features and reduce false positives and negatives. Finally, the extraction of fly outlines was achieved using the contour detection algorithm in the OpenCV library.

### Statistical Tests

Statistical analysis of mating duration assay was described previously^36–38,78,156,157^. More than 50 males (naïve and experienced) were used for mating duration assay. Our experience suggests that the relative mating duration differences between naïve and experienced condition are always consistent; however, both absolute values and the magnitude of the difference in each strain can vary. So, we always include internal controls for each treatment as suggested by previous studies ^168^. Therefore, statistical comparisons were made between groups that were naïvely reared, sexually experienced within each experiment. As mating duration of males showed normal distribution (Kolmogorov-Smirnov tests, *p > 0.05*), we used two-sided Student’s t tests. The mean ± standard error (s.e.m) (*\*\*\*\* = p < 0.0001, *** = p < 0.001, ** = p < 0.01, * = p < 0.05*). To ensure robust statistical analysis, each experimental group included at least 100 male flies (naïve, and sexually experienced). Internal controls were incorporated into every experiment as recommended by Bretman et al. (2011)^168^. Normality of the mating duration data was confirmed using the Kolmogorov-Smirnov test (*p > 0.05*). For group comparisons, two-sided Student’s t-tests were applied to calculate significance levels (*\*\*\*\* = p < 0.0001, *** = p < 0.001, ** = p < 0.01, * = p < 0.05*), Comparisons among three or more groups were performed using one-way ANOVA with Tukey’s HSD post-hoc tests. Two-factor experimental comparisons were performed using two-way ANOVA followed by Sidak’s post-hoc test. while estimation statistics (Claridge-Chang and Assam, 2016) were used to visualize effect sizes, mean differences, and precision, avoiding reliance solely on null hypothesis testing. All analyses, including data plotting, were performed using GraphPad Prism software.

Besides traditional *t*-test for statistical analysis, we added estimation statistics for all MD assays and two group comparing graphs. In short, ‘estimation statistics’ is a simple framework that—while avoiding the pitfalls of significance testing—uses familiar statistical concepts: means, mean differences, and error bars. More importantly, it focuses on the effect size of one’s experiment/intervention, as opposed to significance testing ^169^. In comparison to typical NHST plots, estimation graphics have the following five significant advantages such as (1) avoid false dichotomy, (2) display all observed values (3) visualize estimate precision (4) show mean difference distribution. And most importantly (5) by focusing attention on an effect size, the difference diagram encourages quantitative reasoning about the system under study ^170^. Thus, we conducted a reanalysis of all our two group data sets using both standard *t* tests and estimate statistics. In 2019, the Society for Neuroscience journal eNeuro instituted a policy recommending the use of estimation graphics as the preferred method for data presentation ^171^.

## ACKNOWLEDGMENTS

We thank Dr. Jan A. Veenstra (University of Bordeaux) for sharing *SIFa^PT^-GAL4* driver, Drs. Yuh Nung Jan and Lily Yeh Jan (UCSF, USA) for helpful comments and support on this paper. We are also very appreciative to the colleagues who supplied us with several fly strains: Dr. Wei Zhang (Tsinghua University), Dr. Fang Guo (Zhejiang University), and Dr. Yufeng Pan (Southeast University), Drs. Young-Joon Kim and Sung-Eun Yoon (Korea Drosophila Resource Center, KDRC), and Drs. Kweon Yu and Tae Hoon Ryu (KRIBB). We have greatly benefited from the resources provided by the FlyBase website in our genetic research endeavors. We are grateful for the ongoing efforts of the FlyBase staff in maintaining this comprehensive *Drosophila* database ^172–176^. Furthermore, we extend our sincere thanks to the TsingHua Fly Center (THFC), *Drosophila* Resource and Technology (Shanghai, BCFly) and the broader Chinese fly community, especially Dr. Guo Xuan (Jinzhou Medical University) for his invaluable support, as well as to Qidong Fungene Biotechnology for sharing essential fly strains used in this study. This research was supported by a Startup funds from HIT Center for Life Science to WJK. The funder had no role in study design, data collection and analysis, decision to publish, or preparation of the manuscript.

## AUTHOR CONTRIBUTIONS

**Conceptualization:** Woo Jae Kim.

**Data curation:** Tianmu Zhang, Yanying Sun, Yanan Wei, Wenjing Li, Jie Chen, Shun-Fan Wu, Yutong Song, Kyle Wong, Woo Jae Kim.

**Formal analysis:** Tianmu Zhang, Yanying Sun, Yanan Wei, Wenjing Li, Jie Chen, Shun-Fan Wu, Yutong Song, Kyle Wong, Woo Jae Kim.

**Funding acquisition:** Woo Jae Kim.

**Investigation:** Woo Jae Kim.

**Methodology:** Tianmu Zhang, Yanying Sun, Yanan Wei, Wenjing Li, Woo Jae Kim.

**Project administration:** Woo Jae Kim.

**Resources:** Woo Jae Kim.

**Supervision:** Shanfan Wu, Woo Jae Kim.

**Validation:** Tianmu Zhang, Shanfan Wu, Woo Jae Kim.

**Visualization:** Tianmu Zhang, Woo Jae Kim.

**Writing – original draft:** Woo Jae Kim.

**Writing – review & editing:** Tianmu Zhang, Yanying Sun, Yanan Wei, Woo Jae Kim.

## CONFLICT OF INTERESTS

The authors declare no competing interests.

## STATEMENT OF ANIMAL RESEARCH COMPLIANCE

All animal experiments reported in this manuscript were conducted in compliance with the ARRIVE guidelines and adhered to the U.K. Animals (Scientific Procedures) Act, 1986 and associated guidelines, EU Directive 2010/63/EU for animal experiments, or the National Research Council’s Guide for the Care and Use of Laboratory Animals.

## DECLARATION OF GENERATIVE AI AND AI-ASSISTED TECHNOLOGIES IN THE WRITING PROCESS

During the creation of this work, the author(s) utilized generative AI to rephrase English sentences, verify English grammar, and detect plagiarism, as none of the authors of this paper are native English speakers. After using this tool/service, the author(s) reviewed and edited the content as needed and take(s) full responsibility for the content of the publication.

## DATA AVAILABILITY STATEMENT

Strains are available upon request. The authors affirm that all data necessary for confirming the conclusions of the article are present within the article, figures, and tables.

---

**Figure S1. Long-range SIFa axons innervate the hindgut ampulla and rectal papillae.**

(A) 24-h sucrose intake of males measured by EX-Q assay of knockdown of SIFa in SIFa^PT^-expressing neurons between naïve and experienced male flies on yeast-sugar medium. See the **MATERIALS AND METHODS** for a detailed description of the EX-Q assay used in this study.

(B-D) Fly gut expressing *GAL4^SIFa.PT^* driver together with *UAS-CD4-tdTomato* were immunostained with anti-SIFa (green, for SIFa peptide) and anti-AstA (green, for AstA peptide), anti-tdTomato (red, for SIFa neurons), and anti-BP102 (blue, for axons) antibodies. Boxes indicate the magnified regions of interest presented in the bottom panels. Scale bars represent 100 μm.

(E) Working model summarizing SIFa neuronal mapping.

**Figure S2. SIFaR is not expressed in intrinsic gut cell types.**

(A, B) Analysis of single-cell RNA sequencing datasets from the Fly Cell Atlas (SCope) reveals no detectable expression of the SIFa receptor (SIFaR) in adult gut epithelial lineages, including intestinal stem cells (ISCs; esg□), enteroendocrine cells (EE; pros□), or other gut cell types. These independent datasets corroborate the absence of intrinsic enteric SIFaR expression and support the conclusion that SIFa signaling acts on the gut indirectly via extrinsic neural inputs.

**Figure S3. Peripheral architecture and plasticity of SIFaR projections to gut and reproductive tissues.**

(A) High-magnification imaging of male fly expressing *SIFaR-GAL4* together with *UAS-Denmark, UAS-syteGFP* were immunostained with anti-RFP (red) for dendrites and anti-GFP (green). Scale bars represent 100 μm.

(B) Hindgut and rectum of fly expressing *SIFaR-GAL4* together with *UAS-CD4tdGFP* with anti-GFP (green, for SIFaR), anti-BP102 (red, for axons), and phalloidin (blue, for actin). Projection tracing shows that SIFaR neurons bypass the MHJ and project directly to the hindgut ampulla. Scale bars represent 100 μm in hindgut panels and 50 μm in rectum panels.

(C) Male fly expressing *SIFaR-GAL4* together with *UAS-Denmark*, *UAS-syteGFP* were immunostained with anti-RFP (red) for dendrites and anti-GFP (green). SIFaR neurons in the AG extend robust dendritic and axonal projections into the male reproductive system, despite the absence of direct innervation from central SIFa neurons, indicating an indirect relay pathway for SIFa-dependent modulation of reproductive physiology. Scale bars represent 100 μm in upper panels and 50 μm in lower panels.

(D) Male fly expressing *SIFaR-GAL4* together with *UAS-syteGFP* were immunostained with and anti-GFP (green) between naïve and experienced condition. Scale bars represent 20 μm.

(E-G) Quantification of particle numbers (E), average size (F) and percent of area (G) of SIFaR in rectal papillae. In all plots and statistical tests. Data are presented as mean ± s.e.m. ns = not significant *(p>0.05), *p<0.05, **p<0.01, ***p< 0.001, ****p< 0.0001*. Sample sizes (n) are indicated in the figure panels. For detailed methods, see the **MATERIALS AND METHODS** for a detailed description of the particle analysis used in this study.

(H) Crop of male fly expressing *SIFaR-GAL4* together with *UAS-Denmark, UAS-syteGFP.* SIFaR-positive dendrites (magenta), axons (green), crop axons (cyan). SIFaR-positive neurons in the corpora cardiaca (CC) project extensively to the foregut crop. These projections display overlapping dendritic and axonal architectures similar to those observed at the rectal papillae, identifying the CC as the source of dense SIFaR innervation in the foregut. Scale bars represent 100 μm.

**Figure S4. Functional and anatomical compartmentalization of peripheral SIFaR innervation via OA/TA and AKH neuroendocrine systems.**

(A) Male fly gut expressing *SIFaR^24F06^-GAL4* and *Tdc2-lexA* drivers together with *UAS-mCD8RFP* and *lexAop-mCD8GFP*, was immunostained with anti-GFP (green), anti-DsRed (red) and anti-nc82 (blue) antibodies. Areas outlined by white boxes are enlarged in the right panel. Individual single-channel grayscale inverted displays for SIFaR^24F06^ (red channel), Tdc2 (green channel), and nc82 (blue channel) signals. Boxes indicate the magnified regions of interest presented in the right panels. Scale bars represent 100 μm.

(B) Flies expressing *AKH-lexA* and *SIFaR-GAL4* drivers together with *lexAop-mCD8GFP* and *UAS-mCD8RFP* were immunostained with anti-GFP (green for SIFaR), anti-RFP (red for AKH), and anti-BP102 (blue for CNS axons) antibodies. Scale bars represent 100 μm.

**Figure S5. Functional architecture, tissue-specific receptor deployment, and neurochemical profiling of the octopaminergic-AKH neuroendocrine axis.**

(A) Male flies’ VNC and SOG expressing *AKH-GAL4* together with *UAS-Denmark* and *UAS-syt.eGFP* in naive (upper panels) and exp. (lower panels) conditions were immunostained with anti-GFP (green), and nc82 (magenta) antibodies. Independent single-channel green fluorescence panels (syteGFP) are presented alongside to highlight synaptic structural configurations across social cohorts. Scale bars represent 50 μm.

(B-C) Quantification of relative intensity value for GFP fluorescence in VNC and SOG regions (two-tailed unpaired *t*-test). In all plots and statistical tests. Data are presented as mean ± s.e.m. ns = not significant (*p>0.05*), *\*p<0.05, **p<0.01, ***p< 0.001, ****p< 0.0001*. Sample sizes (n) are indicated in the figure panels.

(D) Mating duration assay evaluating tissue-specific requirement of the AkhR. Genetic knockdown of AkhR within the central nervous system/fan-shaped body (*elav^c155^*) preserves normal SMD responses in experienced males (red data points) relative to naive controls (grey data points). Conversely, restriction of AkhR knockdown to the metabolic fat body storage organ via *Lsp^2.1^-GAL4* completely disrupts the post-mating SMD behavioral switch. In all plots and statistical tests. Data are presented as mean ± s.e.m. ns = not significant *(p>0.05), *p<0.05, **p<0.01, ***p< 0.001, ****p< 0.0001.* Sample sizes (n) are indicated in the figure panels.

(E) Anterograde tracing via the trans-Tango system driven by *AKH-GAL4* along the whole-gut. Specific panels delineate downstream postsynaptic targets (red), intrinsic *AKH-GAL4*-labeled neuroendocrine fibers (green), autofluorescent tissue scaffolding (UV, cyan), and a multi-channel composite showing postsynaptic structures descending towards the foregut crop and hindgut ampulla. Scale bars represent 100 μm.

(F-H) Retrograde neural mapping via the retro-Tango system combined with *AKH-GAL4* to identify upstream presynaptic inputs modulating AKH cells. (F) Strong presynaptic donor structures (red) are localized within the central complex fan-shaped body (FB) of the brain. High-magnification synaptic input profiling within the (G) VNC core and (H) localized CC ring gland anatomy, counterstained with anti-nc82 (blue) to verify central network docking points. Scale bars represent 100 μm in (F), 50 μm in (G), and 20 μm in (H).

(I) Ex-Q assays for GAL4 mediated knockdown of Tdc2 via *Tdc2-RNAi* using the AKH driver.

(J) SMD assays for GAL4 mediated knockdown of Tdc2 via *Tdc2-RNAi* using the AKH driver.

(K) Single-nucleus transcriptomic co-expression mapping derived from the FlyCellAtlas (SCope) dataset, focusing on the corpora cardiaca neuroendocrine cluster. Two-dimensional t-SNE projections demonstrate dense co-localization of the metabolic neuropeptide AKH (Red), the rate-limiting octopaminergic/tyraminergic biosynthetic enzyme Tdc2 (Green), and the receptor SIFaR (Blue) within a unified endocrine cellular population.

**Figure S6. Comparative mapping of central, peripheral, and enteroendocrine AstA pools relative to SIFaR expression domains.**

(A) Male flies expressing *SIFaR-GAL4* and *AstA-lexA* together with *UAS-mCD8RFP* and *lexAop-mCD8GFP*, counterstained for nc82 synaptic organization. Multi-channel merged composite detailing expression signatures across specified transit zones including the crop, CF, CC, midgut R1, R2 and R3. Scale bars represent 100 μm.

(B) Male flies expressing *SIFaR-GAL4* and *AstA-lexA* together with *UAS-mCD8RFP* and *lexAop-mCD8GFP*, counterstained for nc82 synaptic organization. Multi-channel merged composite detailing expression signatures across specified transit zones including the midgut R4, R5, MHJ, and rectum. Scale bars represent 100 μm.

**Figure S7. Social context-dependent plasticity of central SIFa-AstA synapses and brain-specific characterization of receptor lineages.**

(A) Male flies testis expressing *AstA-GAL4^T2A^* together with *UAS-mCD8RFP* were immunostained with anti-RFP (red) antibodies. Scale bars represent 100 μm.

(B) tGRASP assay (resulted in a strong preferential GRASP signal in synaptic regions) for *SIFa^PT^-lexA* and *AstA-GAL4* in brain and VNC of male flies in naive (left panels) and exp. (right panels) conditions. Male flies were dissected after 5 days of growth. Synaptic transmission occurs from *SIFa^PT^-lexA* to *AstA-GAL4.* GFP is pseudo-colored as “fire”. Scale bars represent 100 μm in brain, OL and SOG panels, 50 μm in VNC panels, 25 μm in AG panels.

(C-E) Quantification of relative value for synaptic intensity in OlL, SOG and AG regions (two-tailed unpaired *t*-test). In all plots and statistical tests. Data are presented as mean ± s.e.m. ns = not significant (*p>0.05*), *\*p<0.05, **p<0.01, ***p< 0.001, ****p< 0.0001*. Sample sizes (n) are indicated in the figure panels.

(F) Ex-Q assays for GAL4 mediated knockdown of AstA receptors via *AstA-R1-RNAi* and *AstA-R2-RNAi* using the *elav^c155^* driver together with o*td^FLP^, tub-FRT-GAL80*.

(G) SMD assays for GAL4 mediated knockdown of AstA receptors via *AstA-R1-RNAi* and *AstA-R2-RNAi* using the *elav^c155^* driver together with *otd^FLP^, tub-FRT-GAL80*. (H-I) Baseline expression mapping of receptor driver lines within whole-mount adult brain and VNC complexes using (H) *AstA-R1-GAL4* together with *UAS-mCD8GFP* and (I) *AstA-R2-GAL4* together with *UAS-mCD8GFP*. Scale bars represent 100 μm in brain panels, 50 μm in VNC panels,

**Figure S8. Structural stability of ascending ganglionic VNC synapses and modular dissection of neuronal versus gut-derived AstA pools.**

(A) Confocal evaluation of ascending gut-to-brain feedback networks using the t-GRASP reconstitution system. The representative cross-sections compare the ventral nerve cord (VNC) of adult male flies under naive (left) and sexually experienced (right) social conditions. Synaptic complementation signals (pseudo-colored as “red hot”) reveal physical contact points between ascending SIFaR-expressing interneurons and descending SIFa projections traversing the ganglionic tracks. Scale bars represent 100 μm.

(B) Quantitative analysis of the relative t-GRASP functional connection area (normalized against nc82 neuropil counterstaining) within the VNC neuropils between naive (grey bar) and sexually experienced (red bar) male cohorts. Estimation graphics plot the bootstrap distribution of the difference between means (DBMs). Data are presented as mean ± s.e.m. ns = not significant *(p>0.05), *p<0.05, **p<0.01, ***p< 0.001, ****p< 0.0001.* Sample sizes (n) are indicated in the figure panels.

(C) EX-Q assay mapping the homeostatic requirement of distinct AstA source pools via tissue-specific genetic knockdown AstA with *AstA-RNAi* using *AstA^33^-GAL4* driver and *elav^c155^* driver together with *tsh-GAL80*.

**Figure S9. Expression pattern of SIFa in CNS and hindgut of different insect.** (A-J) Expression pattern of SIFa in CNS and hindgut in (A) *Drosophila virillis*, (B) *Drosophila yakuba*, (C) *Drosophila pseudoobscura*, (D) *Drosophila mojavensis*, (E) *Coccinella septempunctata*, (F) *Acheta domesticus*, (G) *Nilaparvata lugens*, (H) *Messor structor*, (I) *Spodoptera frugiperda*, and (J) *Apis mellifera*. Scale bars represent 100 μm.

## Notes

### Competing Interest Statement

The authors have declared no competing interest.

## REFERENCES

1. Pianka, E. R. Natural Selection of Optimal Reproductive Tactics. Am. Zoo l. 16, 775–784 (1976).

2. Prescott, S. L. & Liberles, S. D. Internal senses of the vagus nerve. Neuron 110, 579–599 (2022).

3. Seoane-Collazo, P. et al. Hypothalamic-autonomic control of energy homeostasis. Endocrine 50, 276–291 (2015).

4. Székely, M. The vagus nerve in thermoregulation and energy metabolism. Auton. Neurosci. 85, 26–38 (2000).

5. Howland, R. H. Vagus Nerve Stimulation. Curr. Behav. Neurosci. Rep. 1, 64–73 (2014).

6. Abdullah, N., Defaye, M. & Altier, C. Neural control of gut homeostasis. Am. J. Physiol.-Gastrointest. Liver Physiol. 319, G718–G732 (2020).

7. Mayer, E. A. Gut feelings: the emerging biology of gut–brain communication. Nat Rev Neurosci 12, 453–466 (2011).

8. Sadaqat, Z., Kaushik, S. & Kain, P. Preclinical Animal Modeling in Medicine. (2022) doi:10.5772/intechopen.96503.

9. Nässel, D. R. & Zandawala, M. Endocrine cybernetics: neuropeptides as molecular switches in behavioural decisions. Open Biol 12, 220174 (2022).

10. Yapici, N. Gut-brain communication in Drosophila melanogaster. Curr. Opin. Neurobiol. 94, 103096 (2025).

11. Scopelliti, A. et al. A Neuronal Relay Mediates a Nutrient Responsive Gut/Fat Body Axis Regulating Energy Homeostasis in Adult Drosophila. Cell Metab 29, 269–284.e10 (2019).

12. Chatterjee, N. & Perrimon, N. What fuels the fly: Energy metabolism in Drosophila and its application to the study of obesity and diabetes. Sci Adv 7, eabg4336 (2021).

13. Droujinine, I. A. & Perrimon, N. Interorgan Communication Pathways in Physiology: Focus on Drosophila. Annu. Rev. Genet. 50, 539–570 (2015).

14. Nässel, D. R. A brief history of insect neuropeptide and peptide hormone research. Cell Tissue Res. 399, 129–159 (2025).

15. Taghert, P. H. & Nitabach, M. N. Peptide Neuromodulation in Invertebrate Model Systems. Neuron 76, 82–97 (2012).

16. Nässel, D. R. & Winther, Å. M. E. Drosophila neuropeptides in regulation of physiology and behavior. Prog Neurobiol 92, 42–104 (2010).

17. Chen, J. et al. Allatostatin A Signalling in Drosophila Regulates Feeding and Sleep and Is Modulated by PDF. PLoS Genet. 12, e1006346 (2016).

18. Hergarden, A. C., Tayler, T. D. & Anderson, D. J. Allatostatin-A neurons inhibit feeding behavior in adult Drosophila. Proc. Natl. Acad. Sci. 109, 3967–3972 (2012).

19. Li, Y. et al. Gut AstA mediates sleep deprivation-induced energy wasting in Drosophila. Cell Discov. 9, 49 (2023).

20. Oh, Y. et al. A glucose-sensing neuron pair regulates insulin and glucagon in Drosophila. Nature 574, 559–564 (2019).

21. Malita, A. et al. A gut-derived hormone suppresses sugar appetite and regulates food choice in Drosophila. Nat Metabolism 1–19 (2022) doi:10.1038/s42255-022-00672-z.

22. Pauls, D. et al. Endocrine signals fine-tune daily activity patterns in Drosophila. Curr. Biol. 31, 4076–4087.e5 (2021).

23. Gáliková, M. et al. Energy Homeostasis Control in Drosophila Adipokinetic Hormone Mutants. Genetics 201, 665–683 (2015).

24. Nässel, D. R., Enell, L. E., Santos, J. G., Wegener, C. & Johard, H. A. A large population of diverse neurons in the Drosophilacentral nervous system expresses short neuropeptide F, suggesting multiple distributed peptide functions. Bmc Neurosci 9, 90 (2008).

25. Semaniuk, U. et al. Drosophila insulin like peptides: from expression to functions – a review. Èntomol. Exp. Appl. 169, 195–208 (2021).

26. Nässel, D. R. & Broeck, J. V. Insulin/IGF signaling in Drosophila and other insects: factors that regulate production, release and post-release action of the insulin-like peptides. Cell. Mol. Life Sci. 73, 271–290 (2016).

27. Ubuka, T. & Tsutsui, K. Evolution of gonadotropin-inhibitory hormone receptor and its ligand. Gen. Comp. Endocrinol. 209, 148–161 (2014).

28. Elphick, M. R. From gonadotropin-inhibitory hormone to SIFamides: Are echinoderm SALMFamides the “missing link” in a bilaterian family of neuropeptides that regulate reproductive processes? Gen. Comp. Endocrinol. 193, 229–233 (2013).

29. Verleyen, P. et al. SIFamide is a highly conserved neuropeptide: a comparative study in different insect species. Biochem Bioph Res Co 320, 334–341 (2004).

30. Arendt, A., Neupert, S., Schendzielorz, J., Predel, R. & Stengl, M. The neuropeptide SIFamide in the brain of three cockroach species. J. Comp. Neurol. 524, 1337–1360 (2016).

31. Veenstra, J. A. The neuropeptide SMYamide, a SIFamide paralog, is expressed by salivary gland innervating neurons in the American cockroach and likely functions as a hormone. bioRxiv 2020.09.17.302331 (2020) doi:10.1101/2020.09.17.302331.

32. Park, S., Sonn, J. Y., Oh, Y., Lim, C. & Choe, J. SIFamide and SIFamide Receptor Define a Novel Neuropeptide Signaling to Promote Sleep in Drosophila. Mol Cells 37, 295–301 (2014).

33. Sellami, A. & Veenstra, J. A. SIFamide acts on fruitless neurons to modulate sexual behavior in Drosophila melanogaster. Peptides 74, 50–56 (2015).

34. Martelli, C. et al. SIFamide Translates Hunger Signals into Appetitive and Feeding Behavior in Drosophila. Cell Reports 20, 464–478 (2017).

35. Terhzaz, S., Rosay, P., Goodwin, S. F. & Veenstra, J. A. The neuropeptide SIFamide modulates sexual behavior in Drosophila. Biochem Bioph Res Co 352, 305– 310 (2007).

36. Zhang, T. et al. Neuropeptide-mediated synaptic plasticity regulates context-dependent mating behaviors in Drosophila. PLOS Biol. 23, e3003330 (2025).

37. Song, Y. et al. SIFa peptidergic neurons orchestrate the internal states and energy balance of male Drosophila melanogaster. PLOS Biol. 23, e3003345 (2025).

38. Zhang, T., Miao, H., Song, Y., Wu, Z. & Kim, W. J. SIFamide-dependent synaptic plasticity in male-specific GABAergic neurons underlies experience-modulated mating behaviors. iScience 28, 113516 (2025).

39. Dreyer, A. P. et al. A circadian output center controlling feeding:fasting rhythms in Drosophila. Plos Genet 15, e1008478 (2019).

40. Kahsai, L. & Winther, Å. M. E. Chemical neuroanatomy of the Drosophila central complex: Distribution of multiple neuropeptides in relation to neurotransmitters. J Comp Neurology 519, 290–315 (2011).

41. Reichl, J., Miller, J. M., Randhawa, H., Castillo, L. M. P. & Akin, O. Four neurons pattern brain-wide developmental activity through neuropeptide signaling. bioRxiv 2025.06.26.661770 (2025) doi:10.1101/2025.06.26.661770.

42. Ayub, M., Hermiz, M., Lange, A. B. & Orchard, I. SIFamide Influences Feeding in the Chagas Disease Vector, Rhodnius prolixus. Front. Neurosci. 14, 134 (2020).

43. Gaskell, W. H. On the Structure, Distribution and Function of the Nerves which innervate the Visceral and Vascular Systems. J. Physiol. 7, 1–80 (1886).

44. Nilsson, S. Comparative anatomy of the autonomic nervous system. Auton. Neurosci. 165, 3–9 (2011).

45. Ottaviani, M. M. & Macefield, V. G. Structure and Functions of the Vagus Nerve in Mammals. Compr. Physiol. 12, 3989–4037 (2022).

46. Yuan, H. & Silberstein, S. D. Vagus Nerve and Vagus Nerve Stimulation, a Comprehensive Review: Part I. Headache: J. Head Face Pain 56, 71–78 (2016).

47. Yuan, H. & Silberstein, S. D. Vagus Nerve and Vagus Nerve Stimulation, a Comprehensive Review: Part II. Headache: J. Head Face Pain 56, 259–266 (2016).

48. Zhang, T. & Kim, W. J. A Master Conductor: How the SIFamide Neuropeptide System Orchestrates Behavioral State in Drosophila. BioEssays 48, e70137 (2026).

49. Yamamoto, D. & Koganezawa, M. Genes and circuits of courtship behaviour in Drosophila males. Nat Rev Neurosci 14, 681–692 (2013).

50. Yamamoto, D., Fujitani, K., Usui, K., Ito, H. & Nakano, Y. From behavior to development: genes for sexual behavior define the neuronal sexual switch in Drosophila. Mech Develop 73, 135–146 (1998).

51. Stockinger, P., Kvitsiani, D., Rotkopf, S., Tirián, L. & Dickson, B. J. Neural Circuitry that Governs Drosophila Male Courtship Behavior. Cell 121, 795–807 (2005).

52. Wasserman, S. A. Multi-layered regulation of courtship behaviour. Nat Cell Biol 2, E145–E146 (2000).

53. Hall, J. C. The Mating of a Fly. Science 264, 1702–1714 (1994).

54. Dauwalder, B. The Roles of Fruitless and Doublesex in the Control of Male Courtship. Int. Rev. Neurobiol. 99, 87–105 (2011).

55. Sun, Y., Zhou, X., Zhang, T., Miao, H. & Kim, W. J. The mating engram: How copulation reshapes the male brain, body, and behavior. Neurosci. Biobehav. Rev. 187, 106776 (2026).

56. Gowaty, P. A., Kim, Y.-K. & Anderson, W. W. No evidence of sexual selection in a repetition of Bateman’s classic study of Drosophila melanogaster. Proc National Acad Sci 109, 11740–11745 (2012).

57. Hoquet, T., Bridges, W. C. & Gowaty, P. A. Bateman’s Data: Inconsistent with “Bateman’s Principles”. Ecol. Evol. 10, 10325–10342 (2019).

58. Douglas, T., Anderson, R. & Saltz, J. B. Limits to male reproductive potential across mating bouts in Drosophila melanogaster. Anim. Behav. 160, 25–33 (2020).

59. Lüpold, S., Manier, M. K., Ala-Honkola, O., Belote, J. M. & Pitnick, S. Male Drosophila melanogaster adjust ejaculate size based on female mating status, fecundity, and age. Behav. Ecol. 22, 184–191 (2011).

60. Sepil, I. et al. Male reproductive aging arises via multifaceted mating-dependent sperm and seminal proteome declines, but is postponable in Drosophila. Proc. Natl. Acad. Sci. 117, 17094–17103 (2020).

61. Ram, K. R. & Wolfner, M. F. Sustained Post-Mating Response in Drosophila melanogaster Requires Multiple Seminal Fluid Proteins. Plos Genet 3, e238 (2007).

62. Gioti, A. et al. Sex peptide of Drosophila melanogaster males is a global regulator of reproductive processes in females. Proc. R. Soc. B: Biol. Sci. 279, 4423–4432 (2012).

63. Kubli, E. Sex-peptides: seminal peptides of the Drosophila male. Cell Mol Life Sci Cmls 60, 1689–1704 (2003).

64. Flatt, T. Survival costs of reproduction in Drosophila. Exp. Gerontol. 46, 369–375 (2011).

65. Bretman, A., Westmancoat, J. D., Gage, M. J. G. & Chapman, T. COSTS AND BENEFITS OF LIFETIME EXPOSURE TO MATING RIVALS IN MALE DROSOPHILA MELANOGASTER. Evolution 67, 2413–2422 (2013).

66. Koppik, M., Ruhmann, H. & Fricke, C. The effect of mating history on male reproductive ageing in Drosophila melanogaster. J. Insect Physiol. 111, 16–24 (2018).

67. Sethi, S. et al. Social Context Enhances Hormonal Modulation of Pheromone Detection in Drosophila. Curr Biol 29, 3887–3898.e4 (2019).

68. Harvanek, Z. M. et al. Perceptive costs of reproduction drive ageing and physiology in male Drosophila. *Nat*. Ecol. Evol. 1, 0152 (2017).

69. Koliada, A. et al. Mating status affects Drosophila lifespan, metabolism and antioxidant system. Comp. Biochem. Physiol. Part A: Mol. Integr. Physiol. 246, 110716 (2020).

70. Lemaître, J. & Gaillard, J. Reproductive senescence: new perspectives in the wild. Biol. Rev. 92, 2182–2199 (2017).

71. Tatar, M., Chien, S. A. & Priest, N. K. Negligible Senescence during Reproductive Dormancy in Drosophila melanogaster. Am. Nat. 158, 248–258 (2001).

72. Sanghvi, K. et al. Reproductive output of old males is limited by seminal fluid, not sperm number. Evol. Lett. 9, 282–291 (2025).

73. Liu, C., Tian, N., Chang, P. & Zhang, W. Mating reconciles fitness and fecundity by switching diet preference in flies. Nat. Commun. 15, 9912 (2024).

74. Ribeiro, C. & Dickson, B. J. Sex Peptide Receptor and Neuronal TOR/S6K Signaling Modulate Nutrient Balancing in Drosophila. Curr. Biol. 20, 1000–1005 (2010).

75. Rushby, H. J., Andrews, Z. B., Piper, M. D. W. & Mirth, C. K. Ageing impairs protein leveraging in a sex-specific manner in Drosophila melanogaster. Anim. Behav. 195, 43–51 (2023).

76. Wu, G. et al. Opposing GPCR signaling programs protein intake setpoint in Drosophila. Cell (2024) doi:10.1016/j.cell.2024.07.047.

77. Wu, Q. et al. Excreta Quantification (EX-Q) for Longitudinal Measurements of Food Intake in Drosophila. iScience 23, 100776 (2020).

78. Lee, S. G. et al. Taste and pheromonal inputs govern the regulation of time investment for mating by sexual experience in male Drosophila melanogaster. PLOS Genet. 19, e1010753 (2023).

79. Wu, Z., Shao, J., Gill, R. & Kim, W. J. Mating duration of male Drosophila melanogaster – A novel genetic model to study interval timing function of human brain. Neurosci. Biobehav. Rev. 176, 106294 (2025).

80. Miao, H., Wu, Z., Wei, Y. & Kim, W. J. CLOCK-dependent pathway in a single pair of LNd neurons instruct circadian-independent interval timing behavior. bioRxiv 2025.09.17.676708 (2025) doi:10.1101/2025.09.17.676708.

81. Zhang, X., Miao, H., Kang, D., Sun, D. & Kim, W. J. Male-specific sNPF peptidergic circuits control energy balance for mating duration through neuron-glia interactions. bioRxiv 2024.10.17.618859 (2024) doi:10.1101/2024.10.17.618859.

82. Sun, D., Zhang, X., Miao, H., Zhang, T. & Kim, W. J. Molecular and Neural Circuit Mechanisms Underlying Sexual Experience-dependent Long-Term Memory in Drosophila. bioRxiv 2024.09.28.615582 (2024) doi:10.1101/2024.09.28.615582.

83. Moshitzky, P. et al. Sex peptide activates juvenile hormone biosynthesis in the Drosophila melanogaster corpus allatum. Arch Insect Biochem 32, 363–374 (1996).

84. Chapman, T. et al. The sex peptide of Drosophila melanogaster: Female post-mating responses analyzed by using RNA interference. Proc National Acad Sci 100, 9923–9928 (2003).

85. Yapici, N., Kim, Y.-J., Ribeiro, C. & Dickson, B. J. A receptor that mediates the post-mating switch in Drosophila reproductive behaviour. Nature 451, 33–37 (2008).

86. Nallasivan, M. P., Singh, D. N., Sahir, M. S. R. & Soller, M. Sex peptide targets distinct higher order processing neurons in the brain to induce the female post-mating response. eLife 13, (2026).

87. Sitnik, J. L., Gligorov, D., Maeda, R. K., Karch, F. & Wolfner, M. F. The Female Post-Mating Response Requires Genes Expressed in the Secondary Cells of the Male Accessory Gland in Drosophila melanogaster. Genetics 202, 1029–1041 (2016).

88. Häsemeyer, M., Yapici, N., Heberlein, U. & Dickson, B. J. Sensory Neurons in the Drosophila Genital Tract Regulate Female Reproductive Behavior. Neuron 61, 511–518 (2009).

89. Yang, C. et al. Control of the Postmating Behavioral Switch in Drosophila Females by Internal Sensory Neurons. Neuron 61, 519–526 (2009).

90. Apger-McGlaughon, J. & Wolfner, M. F. Post-mating change in excretion by mated Drosophila melanogaster females is a long-term response that depends on sex peptide and sperm. J. Insect Physiol. 59, 1024–1030 (2013).

91. White, M. A., Bonfini, A., Wolfner, M. F. & Buchon, N. Drosophila melanogaster sex peptide regulates mated female midgut morphology and physiology. Proc. Natl. Acad. Sci. 118, e2018112118 (2021).

92. Cohen, E., Sawyer, J. K., Peterson, N. G., Dow, J. A. T. & Fox, D. T. Physiology, Development, and Disease Modeling in the Drosophila Excretory System. Genetics 214, 235–264 (2020).

93. Apidianakis, Y. & Rahme, L. G. Drosophila melanogaster as a model for human intestinal infection and pathology. Dis. Model. Mech. 4, 21–30 (2010).

94. Jørgensen, L. M., Hauser, F., Cazzamali, G., Williamson, M. & Grimmelikhuijzen, C. J. P. Molecular identification of the first SIFamide receptor. Biochem. Biophys. Res. Commun. 340, 696–701 (2006).

95. Deng, B. et al. Chemoconnectomics: Mapping Chemical Transmission in Drosophila. Neuron 101, 876–893.e4 (2019).

96. Li, H. et al. Fly Cell Atlas: A single-nucleus transcriptomic atlas of the adult fruit fly. Science 375, eabk2432 (2022).

97. Ma, L., Wang, H.-B. & Hashimoto, K. The vagus nerve: An old but new player in brain–body communication. *Brain, Behav.*, Immun. 124, 28–39 (2025).

98. Nicolaï, L. J. J. et al. Genetically encoded dendritic marker sheds light on neuronal connectivity in Drosophila. Proc National Acad Sci 107, 20553–20558 (2010).

99. Wigglesworth, V. B. The Utilization of Reserve Substances in Drosophila During Flight. J. Exp. Biol. 26, 150–163 (1949).

100. Kim, S. K. & Rulifson, E. J. Conserved mechanisms of glucose sensing and regulation by Drosophila corpora cardiaca cells. Nature 431, 316–320 (2004).

101. Bharucha, K. N., Tarr, P. & Zipursky, S. L. A glucagon-like endocrine pathway in Drosophila modulates both lipid and carbohydrate homeostasis. J. Exp. Biol. 211, 3103–3110 (2008).

102. Lee, G. & Park, J. H. Hemolymph sugar homeostasis and starvation-induced hyperactivity affected by genetic manipulations of the adipokinetic hormone-encoding gene in Drosophila melanogaster. Genetics 167, 311–23 (2004).

103. Shearin, H. K., Quinn, C. D., Mackin, R. D., Macdonald, I. S. & Stowers, R. S. t-GRASP, a targeted GRASP for assessing neuronal connectivity. J Neurosci Meth 306, 94–102 (2018).

104. Roeder, T. TYRAMINE AND OCTOPAMINE: Ruling Behavior and Metabolism. Annu Rev Entomol 50, 447–477 (2005).

105. Song, Y. et al. Peptidergic neurons with extensive branching orchestrate the internal states and energy balance of male Drosophila melanogaster. bioRxiv 2024.06.04.597277 (2024) doi:10.1101/2024.06.04.597277.

106. Talay, M. et al. Transsynaptic Mapping of Second-Order Taste Neurons in Flies by trans-Tango. Neuron 96, 783–795.e4 (2017).

107. Sorkaç, A. et al. retro-Tango enables versatile retrograde circuit tracing in Drosophila. eLife 12, e85041 (2023).

108. Zhang, Y., Zhou, Y., Zhang, X., Wang, L. & Zhong, Y. Clock neurons gate memory extinction in Drosophila. Curr. Biol. 31, 1337–1343.e4 (2021).

109. Huang, H., Possidente, D. R. & Vecsey, C. G. Optogenetic activation of SIFamide (SIFa) neurons induces a complex sleep-promoting effect in the fruit fly Drosophila melanogaster. Physiol Behav 239, 113507 (2021).

110. Rosenzweig, M. et al. The Drosophila ortholog of vertebrate TRPA1 regulates thermotaxis. Genes Dev. 19, 419–424 (2005).

111. Kang, K. et al. Analysis of Drosophila TRPA1 reveals an ancient origin for human chemical nociception. Nature 464, 597–600 (2010).

112. Kang, K. et al. Modulation of TRPA1 thermal sensitivity enables sensory discrimination in Drosophila. Nature 481, 76–80 (2012).

113. Hentze, J. L., Carlsson, M. A., Kondo, S., Nässel, D. R. & Rewitz, K. F. The Neuropeptide Allatostatin A Regulates Metabolism and Feeding Decisions in Drosophila. Sci Rep-uk 5, 11680 (2015).

114. Deveci, D., Martin, F. A., Leopold, P. & Romero, N. M. AstA Signaling Functions as an Evolutionary Conserved Mechanism Timing Juvenile to Adult Transition. Curr. Biol. 29, 813–822.e4 (2019).

115. Veenstra, J. A., Agricola, H.-J. & Sellami, A. Regulatory peptides in fruit fly midgut. Cell Tissue Res. 334, 499–516 (2008).

116. Kendroud, S. et al. Structure and development of the subesophageal zone of the Drosophila brain. II. Sensory compartments. J. Comp. Neurol. 526, 33–58 (2018).

117. Miroschnikow, A., Schlegel, P. & Pankratz, M. J. Making Feeding Decisions in the Drosophila Nervous System. Curr. Biol. 30, R831–R840 (2020).

118. Schoofs, A., Hückesfeld, S. & Pankratz, M. J. Serotonergic network in the subesophageal zone modulates the motor pattern for food intake in Drosophila. J. Insect Physiol. 106, 36–46 (2018).

119. Pan, Y. et al. Differential roles of the fan-shaped body and the ellipsoid body in Drosophila visual pattern memory. Learn Memory 16, 289–295 (2009).

120. Liu, Q., Liu, S., Kodama, L., Driscoll, M. R. & Wu, M. N. Two Dopaminergic Neurons Signal to the Dorsal Fan-Shaped Body to Promote Wakefulness in Drosophila. Curr. Biol. 22, 2114–2123 (2012).

121. Currier, T. A., Matheson, A. M. & Nagel, K. I. Encoding and control of orientation to airflow by a set of Drosophila fan-shaped body neurons. eLife 9, e61510 (2020).

122. Jones, J. D. et al. The dorsal fan-shaped body is a neurochemically heterogeneous sleep-regulating center in Drosophila. PLOS Biol. 23, e3003014 (2025).

123. Kato, Y. S., Tomita, J. & Kume, K. Interneurons of fan-shaped body promote arousal in Drosophila. PLOS ONE 17, e0277918 (2022).

124. De, J., Wu, M., Lambatan, V., Hua, Y. & Joiner, W. J. Re-examining the role of the dorsal fan-shaped body in promoting sleep in Drosophila. Curr. Biol. 33, 3660–3668.e4 (2023).

125. Hu, W. et al. Fan-Shaped Body Neurons in the Drosophila Brain Regulate Both Innate and Conditioned Nociceptive Avoidance. Cell Rep. 24, 1573–1584 (2018).

126. Wong, K., Schweizer, J., Nguyen, K.-N. H., Atieh, S. & Kim, W. J. Neuropeptide relay between SIFa signaling controls the experience-dependent mating duration of male Drosophila. bioRxiv 819045 (2019) doi:10.1101/819045.

127. Landayan, D., Feldman, D. S. & Wolf, F. W. Satiation state-dependent dopaminergic control of foraging in Drosophila. Sci. Rep. 8, 5777 (2018).

128. Frighetto, G., Zordan, M. A., Castiello, U. & Megighian, A. Action-based attention in Drosophila melanogaster. J. Neurophysiol. 121, 2428–2432 (2019).

129. Landayan, D., Wang, B. P., Zhou, J. & Wolf, F. W. Thirst interneurons that promote water seeking and limit feeding behavior in Drosophila. eLife 10, e66286 (2021).

130. Finkelstein, R. & Boncinelli, E. From fly head to mammalian forebrain: the story of otd and Otx. Trends Genet. 10, 310–315 (1994).

131. Theodosiou, N. A. & Xu, T. Use of FLP/FRT System to StudyDrosophilaDevelopment. Methods 14, 355–365 (1998).

132. Zaspel, J. M. The Insects: An Outline of Entomology, 5th Edition. Am. Èntomol. 62, 129–130 (2016).

133. Ryu, S. et al. Transcriptional regulation of neuropeptide receptors underlies context dependent adaptation in Drosophila melanogaster. FEBS Open Bio 16, 90– 115 (2025).

134. Moran, T. H. Cholecystokinin and satiety: current perspectives. Nutrition 16, 858–865 (2000).

135. Roeder, T. Octopamine in invertebrates. Prog. Neurobiol. 59, 533–561 (1999).

136. Grönke, S. et al. Dual Lipolytic Control of Body Fat Storage and Mobilization in Drosophila. PLoS Biol. 5, e137 (2007).

137. Parker, G. A. & Pizzari, T. Sperm competition and ejaculate economics. Biol Rev 85, 897–934 (2010).

138. Yapici, N., Kim, Y.-J., Ribeiro, C. & Dickson, B. J. A receptor that mediates the post-mating switch in Drosophila reproductive behaviour. Nature 451, 33–37 (2008).

139. Sano, H. et al. The Nutrient-Responsive Hormone CCHamide-2 Controls Growth by Regulating Insulin-like Peptides in the Brain of Drosophila melanogaster. PLoS Genet. 11, e1005209 (2015).

140. Hartenstein, V. Development of the insect stomatogastric nervous system. Trends Neurosci. 20, 421–427 (1997).

141. Hartenstein, V. The neuroendocrine system of invertebrates: a developmental and evolutionary perspective. J. Endocrinol. 190, 555–570 (2006).

142. Kaneto, A., Miki, E. & Kosaka, K. Effects of Vagal Stimulation on Glucagon and Insulin Secretion. Endocrinology 95, 1005–1010 (1974).

143. Andresen, M. C. & Paton, J. F. R. Central Regulation of Autonomic Functions. 23–46 (2011) doi:10.1093/acprof:oso/9780195306637.003.0002.

144. Bajar, B. T. et al. A discrete neuronal population coordinates brain-wide developmental activity. Nature 602, 639–646 (2022).

145. Seaver, B. Honey bee social immunity and Colony Collapse Disorder. J. Apic. Res. 50, 87–88 (2011).

146. Morfin, N., Anguiano-Baez, R. & Guzman-Novoa, E. Honey Bee (Apis mellifera) Immunity. Vet. Clin. North Am.: Food Anim. Pr. 37, 521–533 (2021).

147. Vilcinskas, A. Evolutionary plasticity of insect immunity. J. Insect Physiol. 59, 123–129 (2013).

148. Cremer, S., Armitage, S. A. O. & Schmid-Hempel, P. Social Immunity. Curr Biol 17, R693–R702 (2007).

149. Conroy, T. E. & Holman, L. Social immunity in the honey bee: do immune-challenged workers enter enforced or self-imposed exile? Behav. Ecol. Sociobiol. 76, 32 (2022).

150. Maori, E. et al. A Transmissible RNA Pathway in Honey Bees. Cell Reports 27, 1949–1959.e6 (2019).

151. Hyun, U. & Sohn, J.-W. Autonomic control of energy balance and glucose homeostasis. Exp. Mol. Med. 54, 370–376 (2022).

152. Myers, M. G., Affinati, A. H., Richardson, N. & Schwartz, M. W. Central nervous system regulation of organismal energy and glucose homeostasis. Nat. Metab. 3, 737–750 (2021).

153. Zhao, X. & Karpac, J. The Drosophila midgut and the systemic coordination of lipid-dependent energy homeostasis. Curr. Opin. Insect Sci. 41, 100–105 (2020).

154. Kühnlein, R. P. Sensory and Metabolic Control of Energy Balance. Results Probl. Cell Differ. 52, 159–173 (2010).

155. Lee, S. G. et al. Taste and pheromonal inputs govern the regulation of time investment for mating by sexual experience in male Drosophila melanogaster. PLOS Genet. 19, e1010753 (2023).

156. Kim, W. J., Jan, L. Y. & Jan, Y. N. Contribution of visual and circadian neural circuits to memory for prolonged mating induced by rivals. Nat Neurosci 15, 876–883 (2012).

157. Kim, W. J., Jan, L. Y. & Jan, Y. N. A PDF/NPF Neuropeptide Signaling Circuitry of Male Drosophila melanogaster Controls Rival-Induced Prolonged Mating. Neuron 80, 1190–1205 (2013).

158. Song, Y. et al. Simple Methods to Acutely Measure Multiple Timing Metrics among Sexual Repertoire of Male Drosophila. eLife (2026) doi:10.7554/elife.109742.1.

159. Song, Y. et al. Simple Methods to Acutely Measure Multiple Timing Metrics among Sexual Repertoire of Male Drosophila. bioRxiv 2025.11.11.687834 (2025) doi:10.1101/2025.11.11.687834.

160. Parks, A. L. et al. Systematic generation of high-resolution deletion coverage of the Drosophila melanogaster genome. Nat. Genet. 36, 288–292 (2004).

161. Huang, Y., Kwan, A. & Kim, W. J. Y chromosome genes interplay with interval timing in regulating mating duration of male Drosophila melanogaster. Gene Rep. 101999 (2024) doi:10.1016/j.genrep.2024.101999.

162. Zhang, T., Zhang, X., Sun, D. & Kim, W. J. Exploring the Asymmetric Body’s Influence on Interval Timing Behaviors of Drosophila melanogaster. Behav. Genet. 1– 10 (2024) doi:10.1007/s10519-024-10193-y.

163. Feinberg, E. H. et al. GFP Reconstitution Across Synaptic Partners (GRASP) Defines Cell Contacts and Synapses in Living Nervous Systems. Neuron 57, 353–363 (2008).

164. Kayser, M. S., Yue, Z. & Sehgal, A. A Critical Period of Sleep for Development of Courtship Circuitry and Behavior in *Drosophila*. Science 344, 269–274 (2014).

165. Li, H. et al. Fly Cell Atlas: A single-nucleus transcriptomic atlas of the adult fruit fly. Science 375, eabk2432 (2022).

166. Wu, Q. et al. Excreta Quantification (EX-Q) for Longitudinal Measurements of Food Intake in Drosophila. iScience 23, 100776 (2020).

167. Wei, Y. et al. Investigating the immunomodulatory effects of honeybee venom peptide apamin in Drosophila platforms. Infect. Immun. 93, e00131–25 (2025).

168. Bretman, A., Westmancoat, J. D., Gage, M. J. G. & Chapman, T. Males Use Multiple, Redundant Cues to Detect Mating Rivals. Curr. Biol. 21, 617–622 (2011).

169. Claridge-Chang, A. & Assam, P. N. Estimation statistics should replace significance testing. Nat. Methods 13, 108–109 (2016).

170. Ho, J., Tumkaya, T., Aryal, S., Choi, H. & Claridge-Chang, A. Moving beyond P values: data analysis with estimation graphics. Nat. Methods 16, 565–566 (2019).

171. Bernard, C. Estimation Statistics, One Year Later. eNeuro 8, ENEURO.0091-21.2021 (2021).

172. Gramates, L. S. et al. FlyBase: a guided tour of highlighted features. Genetics 220, iyac035 (2022).

173. Attrill, H. et al. FlyBase: establishing a Gene Group resource for Drosophila melanogaster. Nucleic Acids Res. 44, D786–D792 (2016).

174. Öztürk-Çolak, A. et al. FlyBase: updates to the Drosophila genes and genomes database. GENETICS 227, iyad211 (2024).

175. Pierre, S. S. & McQuilton, P. Inside FlyBase: Biocuration as a career. Fly 3, 112–114 (2009).

176. Jenkins, V. K., Larkin, A., Thurmond, J. & Consortium, F. Using FlyBase: A Database of Drosophila Genes and Genetics. Methods Mol. Biol. (Clifton, NJ) 2540, 1–34 (2022).

