## Supplementary figures and images for "A Conserved SIFamide Circuit Functions as the Insect Vagus Nerve"

### Figure S1

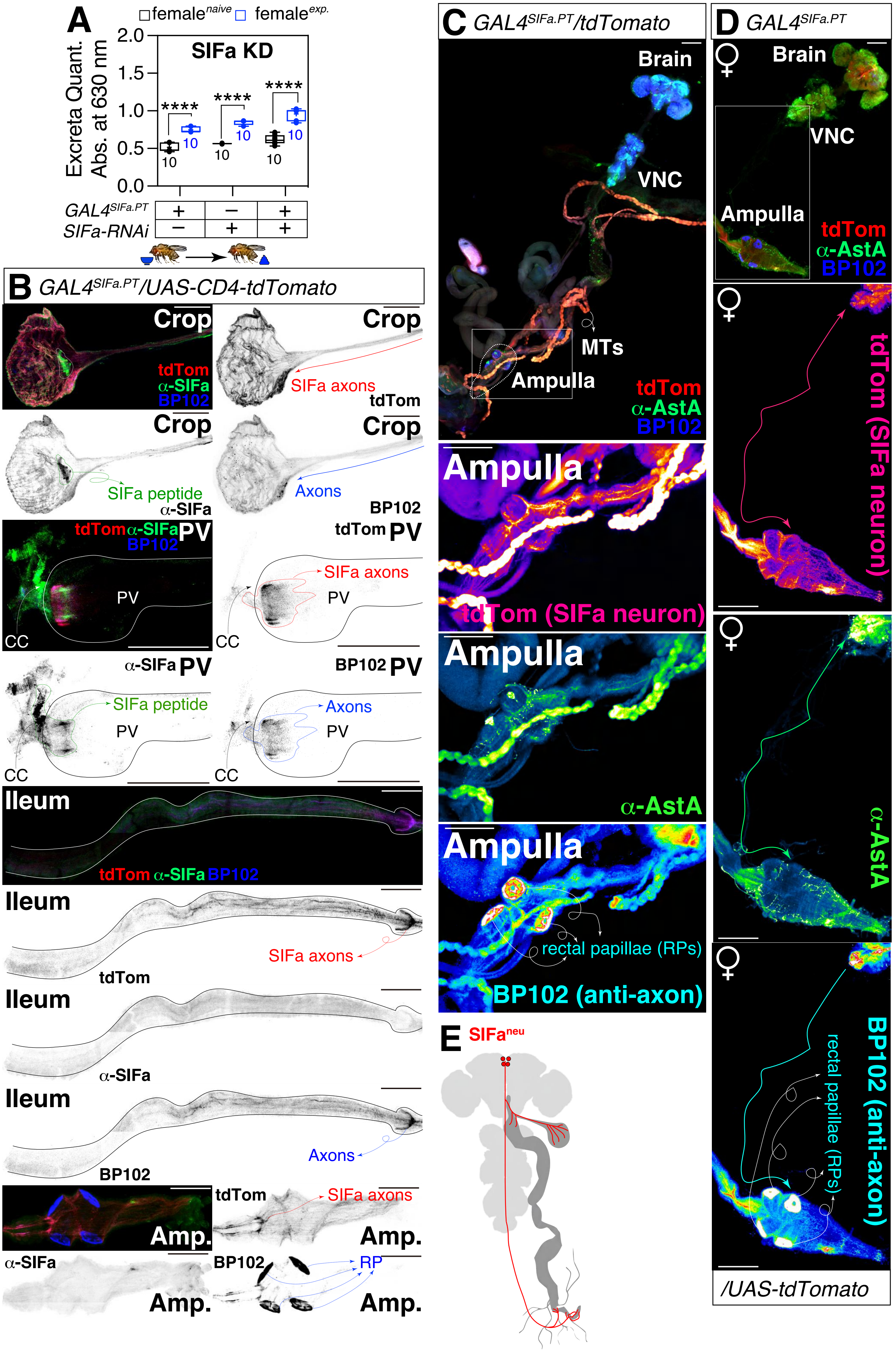


**Vagus-SIFa-Fig.S1**

### Figure S2

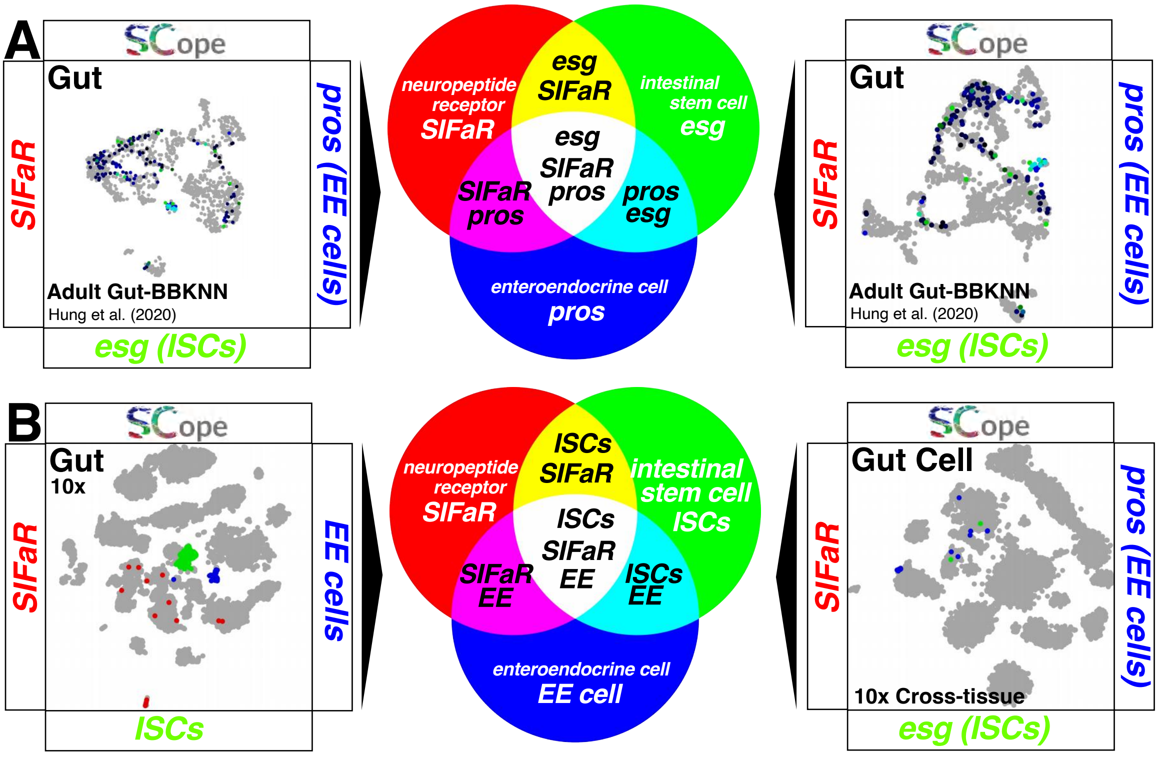


**Vagus-SIFa-Fig.S2**

### Figure S3

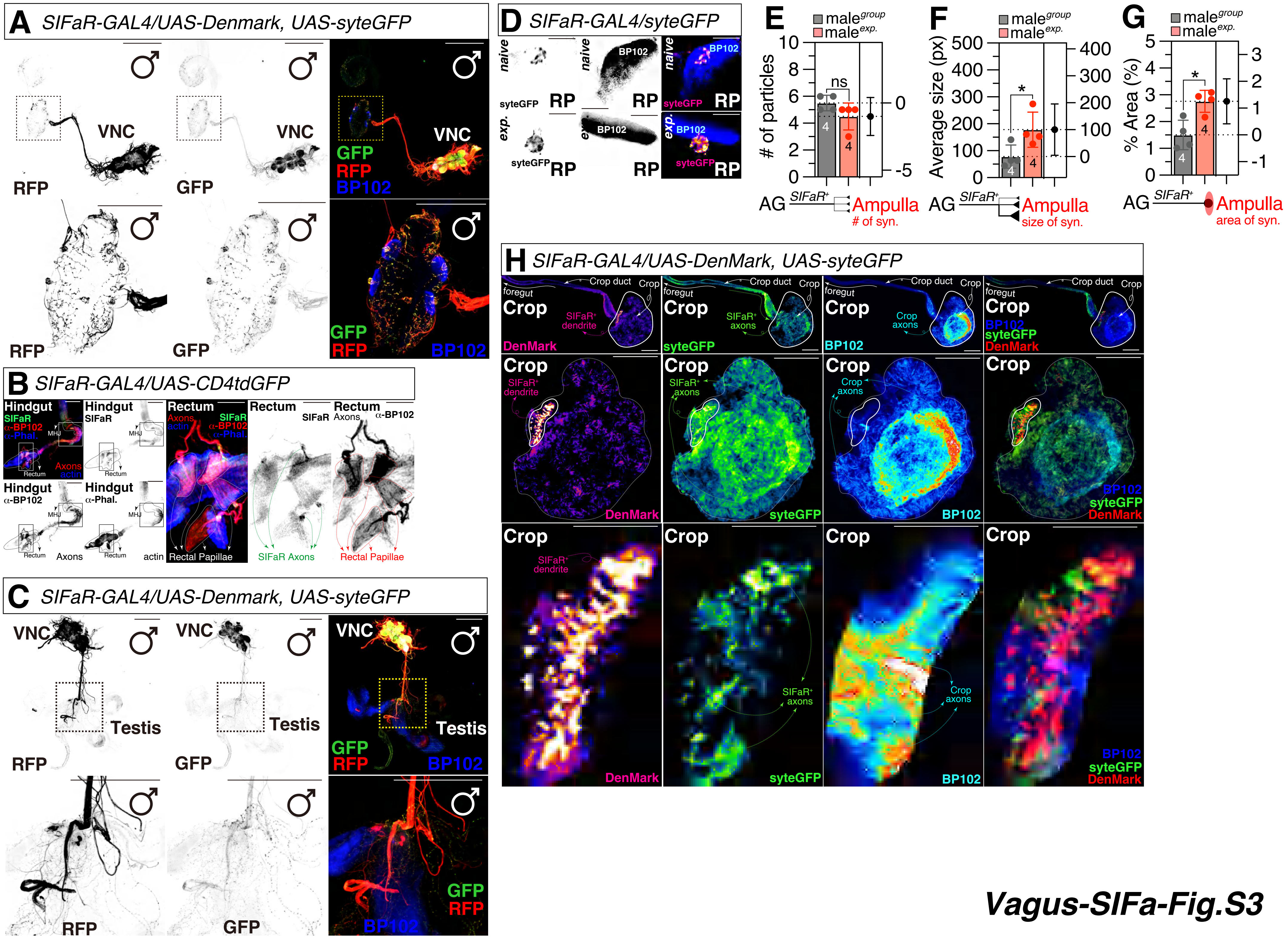

### Figure S4

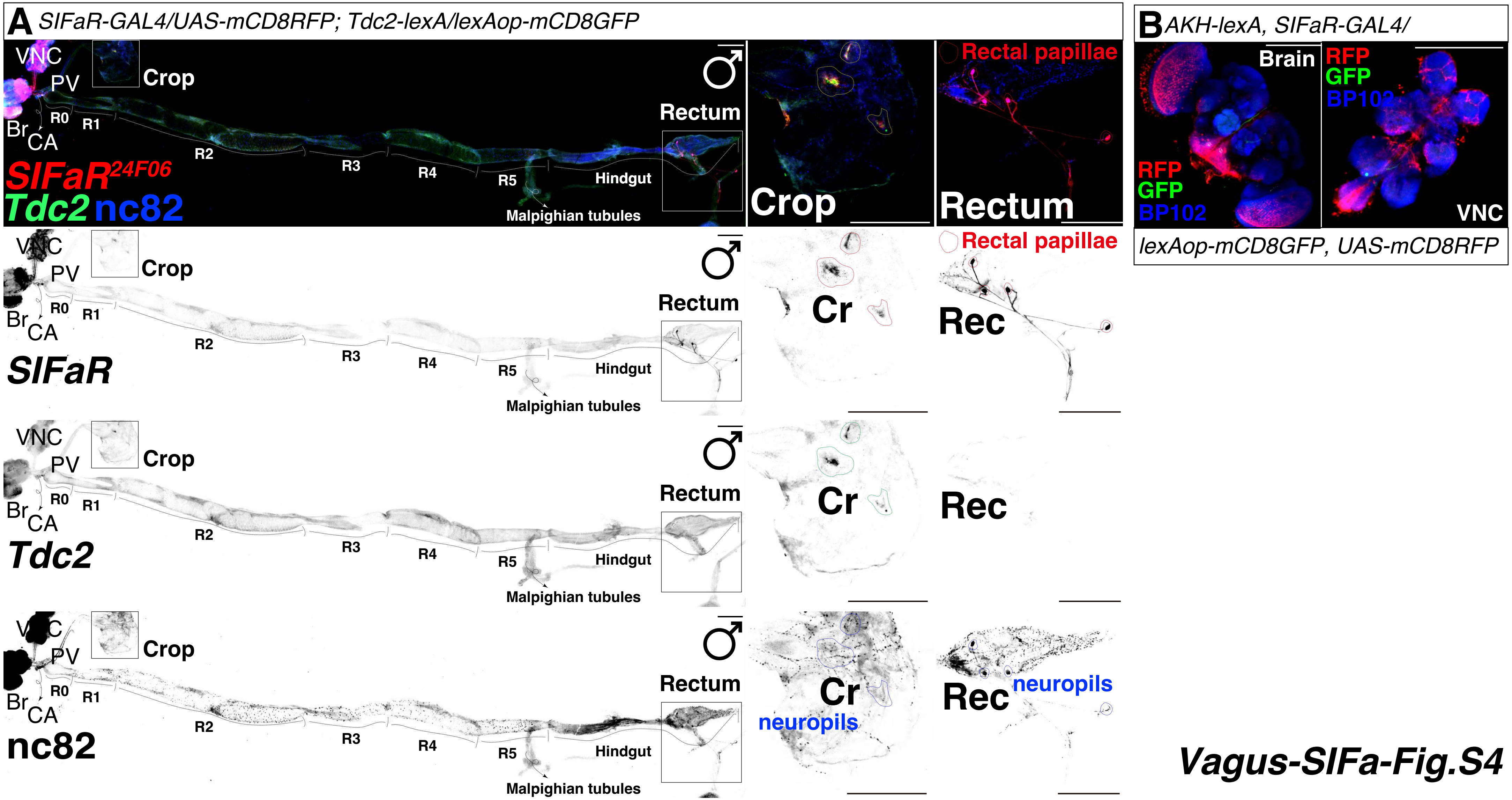


*Vagus-SIFa-Fig.S4*

### Figure S5

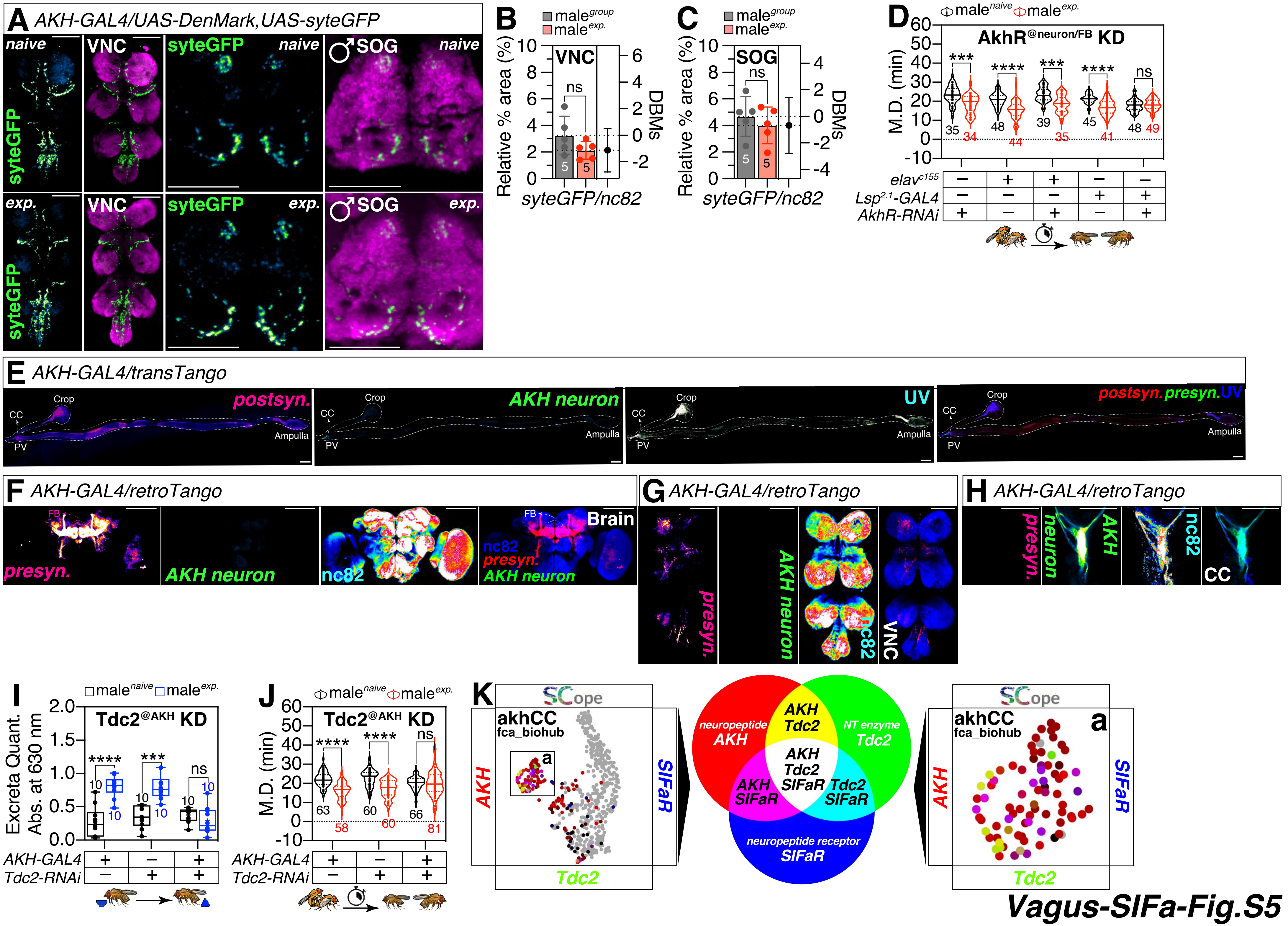

### Figure S6

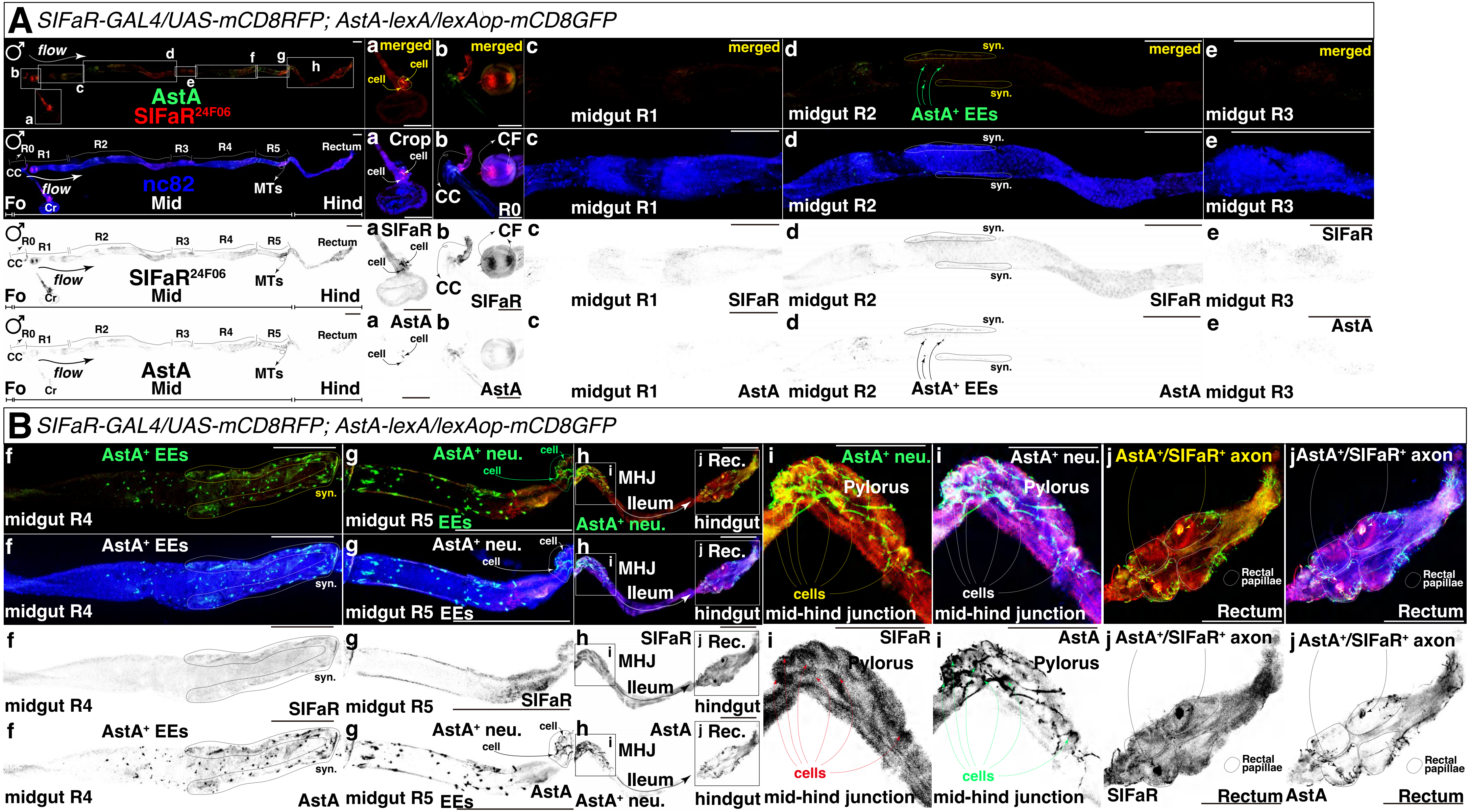


Vagus-SIFa-Fig.S6

### Figure S7

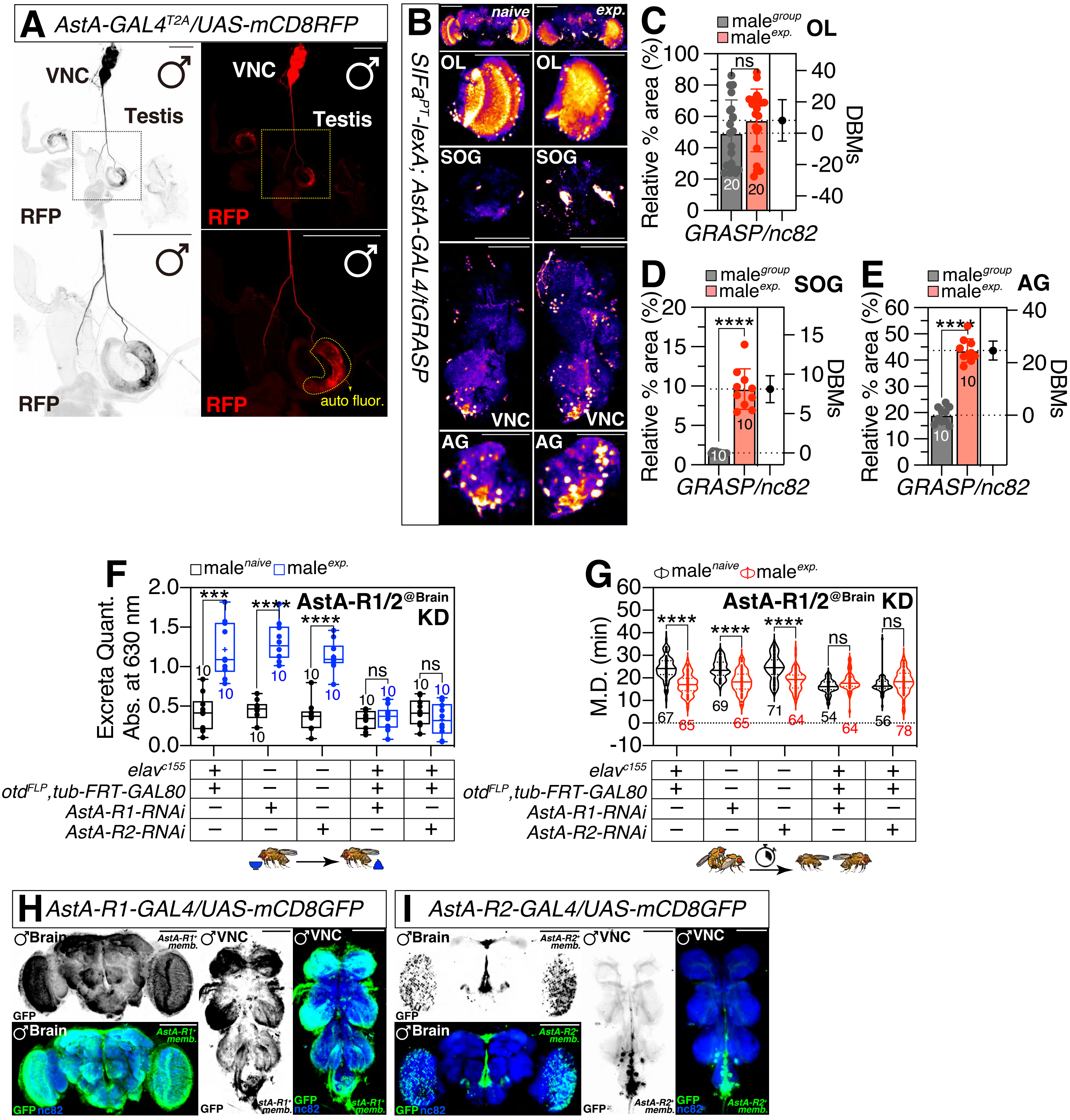


Vagus-SIFa-Fig.S7

### Figure S8

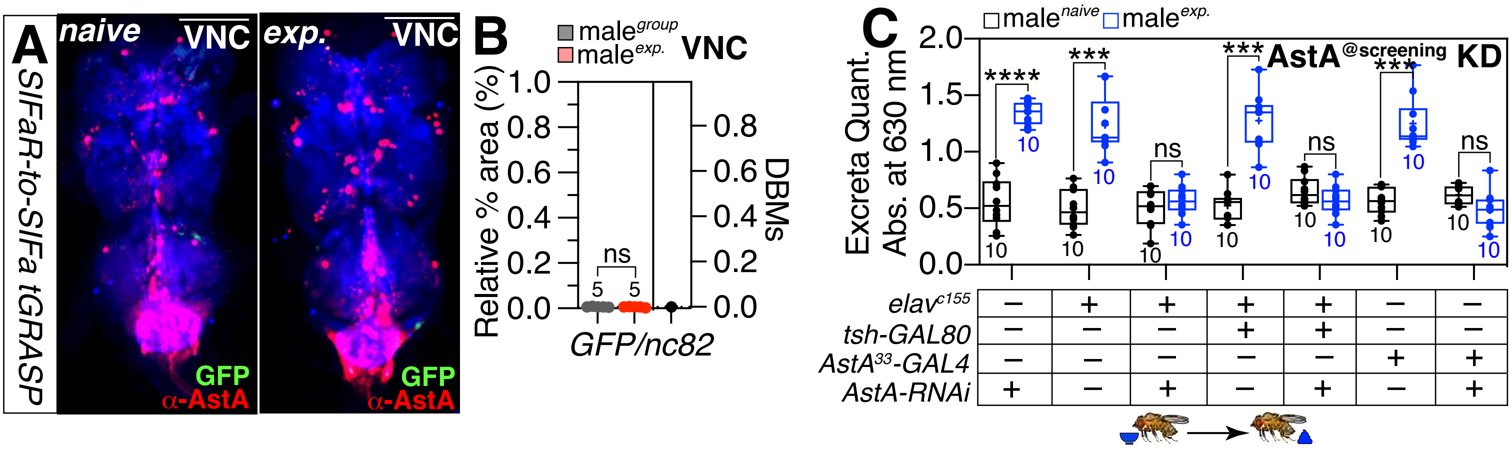


Vagus-SIFa-Fig.S8

### Figure S9

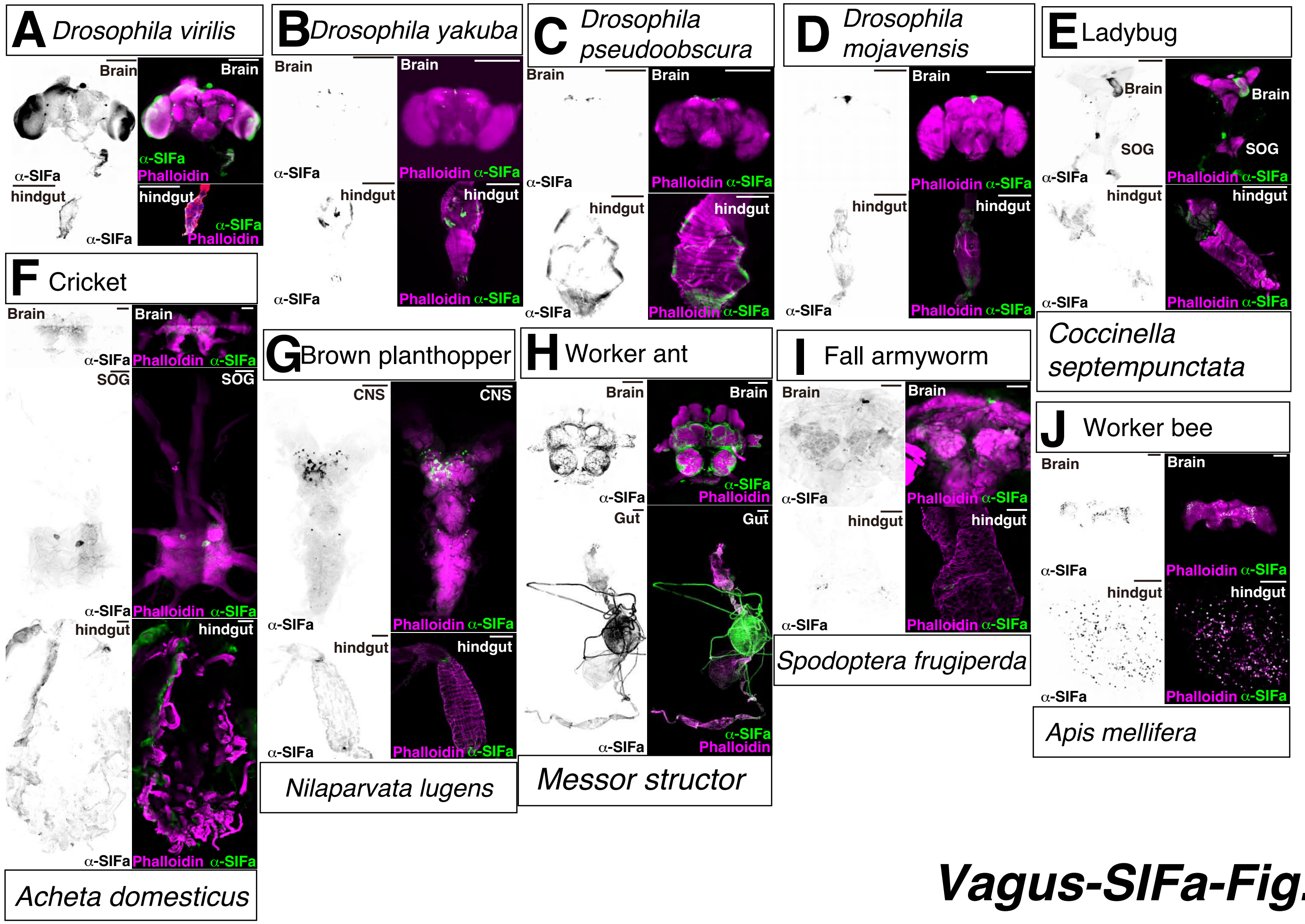


**Vagus-SIFa-Fig.S9**
